# Thymidine Phosphorylase Promotes the Formation of Abdominal Aortic Aneurysm in Mice Fed a Western Diet

**DOI:** 10.1101/2024.02.27.582208

**Authors:** Liang Hong, Hong Yue, Dunpeng Cai, Autumn DeHart, Gretel Toloza-Alvarez, Lili Du, Xianwu Zhou, Xiaoping Fan, Huanlei Huang, Shiyou Chen, Shaik O. Rahaman, Jian Zhuang, Wei Li

## Abstract

**Aims:** The precise molecular drivers of abdominal aortic aneurysm (AAA) remain unclear. Thymidine phosphorylase (TYMP) contributes to increased platelet activation, thrombosis, and inflammation, all of which are key factors in AAA development. Additionally, TYMP suppresses the proliferation of vascular smooth muscle cells (VSMCs), which are central to the development and progression of AAA. We hypothesize that TYMP plays a key role in AAA development.

**Methods and Results:** We conducted a histological study using human AAA samples and normal abdominal aortas, revealing heightened levels of TYMP in human AAA vessel walls. To validate this observation, we utilized an Ang II perfusion-induced AAA model in wild-type C57BL/6J (WT) and *Tymp^−/−^*mice, feeding them a Western diet (TD.88137) starting from 4 weeks of age. We found that *Tymp^−/−^*mice were protected from Ang II perfusion-induced AAA formation. Furthermore, by using TYMP-expressing VSMCs as well as primarily cultured VSMCs from WT and *Tymp^−/−^* mice, we elucidated the essential role of TYMP in regulating MMP2 expression and activation. TYMP deficiency or inhibition by tipiracil, a selective TYMP inhibitor, led to reduced MMP2 production, release, and activation in VSMCs. Additionally, TYMP was found to promote pro-inflammatory cytokine expression systemically, and its absence attenuates TNF-α-stimulated activation of MMP2 and AKT. By co-culturing VSMCs and platelets, we observed that TYMP-deficient platelets had a reduced inhibitory effect on VSMC proliferation compared to WT platelets. Moreover, TYMP appeared to enhance the expression of activated TGFβ1 in cultured VSMCs in vitro and in human AAA vessel walls in vivo. TYMP also boosted the activation of thrombospondin-1 type 1 repeat domain-enhanced TGFβ1 signaling, resulting in increased connective tissue growth factor production.

**Conclusion:** Our findings collectively demonstrated that TYMP serves as a novel regulatory force in vascular biology, exerting influence over VSMC functionality and inflammatory responses that promote the development of AAA.

**Translational Perspective:** Thymidine phosphorylase (TYMP) is increased in the vessel walls of patients with abdominal aortic aneurysm (AAA), and TYMP deficiency in mice reduces the incidence of AAA, suggesting that TYMP plays a crucial role in AAA development. This could be attributed to TYMP’s role in enhancing systemic inflammation and thrombosis, inhibiting vascular smooth muscle cell function, increasing the activation of matrix metalloproteinase and AKT, as well as enhancing the expression of TGFβ1 and connective tissue growth factor. Tipiracil is an FDA-approved drug known to inhibit TYMP-enhanced thrombosis. Targeting TYMP with tipiracil could represent a promising new therapeutic strategy for AAA development.

## 1. Introduction

Abdominal aortic aneurysm (AAA) is a serious vascular condition characterized by a localized dilation of the abdominal aorta (AA) and poses a significant risk, particularly among older men. It is more prevalent in Western nations, presenting a substantial challenge to healthcare systems^1, 2^. The rupture of AAA is frequently fatal in asymptomatic patients, contributing to at least 4.5 deaths per 1,000 individuals, which underscores the severity of this condition^3^. The only effective intervention for symptomatic AAA patients is surgery, through either traditional surgical procedures or endovascular grafting and stenting. However, these procedures are invasive, associated with numerous complications, and require post-surgery supportive treatments. Clinical studies have also indicated limited long-term benefits from these surgical interventions^4^.

Several factors influence the risk of AAA, including age, male gender, hypertension, smoking, obesity, hyperlipidemia, and genetic disorders^4^. Additionally, a Western diet (WD), enhancing systemic low-grade inflammation, has been identified as a contributor to AAA development^5^. The manifestation of AAA involves changes in the structure and function of vascular smooth muscle cells (VSMCs), the extracellular matrix, elastin, and the accumulation of pro-inflammatory agents and cytokines. The progression of AAA can span decades and may remain asymptomatic. Currently, non-surgical treatments for AAA are still in the exploratory phase and lack firm establishment. Furthermore, once an AAA has developed, there are no proven medical therapies available to halt its progression. Therefore, urgent research is needed to explore new cellular and molecular mechanisms that could potentially lead to effective treatments.

The onset of AAA involves multiple genetic factors, including the low-density lipoprotein (LDL) receptor, matrix metalloproteinases (MMPs), particularly MMP2 and MMP9, transforming growth factor-beta1 (TGF-β1), and angiotensin II (Ang II)^6^. These factors are implicated in the weakening of the aortic wall, leading to the loss of VSMCs and increased degradation of the extracellular matrix in the tunica media, crucial aspects of AAA pathogenesis. Thymidine phosphorylase (TYMP), also known as platelet-derived endothelial cell growth factor, is expressed by the endothelial cells, platelets, and certain inflammatory cells. TYMP plays a significant role in platelet activation and thrombosis^7–9^, enhancing chemotaxis in endothelial cells while inhibiting VSMC proliferation^10–12^. Our previous report has hinted at a potential association between TYMP and MMP2/9 in the angiogenic processes^13^. However, the specific role of TYMP in the AAA environment remains unexplored. This study aims to investigate the hypothesis that TYMP, by enhancing MMPs expression and activity in VSMCs and intensifying systemic inflammation, contributes to the reduction of aortic wall integrity, thereby promoting AAA development and progression.

## 2. Materials and Methods

### 2.1. Animals

This study utilized wildtype (WT) and *Tymp^−/−^* mice^9^ in C57BL/6J background, with 26 mice in each genotype. The mice were provided *ad libitum* access to a standard laboratory rodent diet and water. Given the significant influence of female sex hormones on the development of vascular diseases^14^, this study focused on male mice aged 4 to 16 weeks to investigate the proposed concept. Ethical approval for the animal studies was obtained from the Institutional Animal Care and Use Committee (IACUC) of Marshall University (IACUC#: 1033528, PI: Wei Li).

### 2.2. Establishment of AAA model and vascular ultrasonography

We adopted the AAA model as mentioned by Dr. Daugherty et al.^15^ with some modifications. Mice were fed WD (TD88137, Envigo) starting at 4 weeks of age for 8 weeks. Subsequently, mice received implantation of an Ang II-containing osmotic mini pump (Alzet®, Model 2004), which delivered Ang II (Alomone labs, Cat# GPA-100) to mice at a dose of 1 µg/kg/min for 4 weeks. For this procedure, mice were anesthetized with 5% isoflurane mixed with 100% O_2_ at a flow rate of 1 L/min. The depth of anesthesia was confirmed by toe pinching and maintained with 1.5 - 2% isoflurane with 100% O_2_. The hair on the dorsal region of the neck was removed, and the skin was sanitized with Povidone Iodine Wipes followed by Alcohol Wipes. The mouse was covered with a sterilized surgical sheet with a 2 x 2 cm hole in the center to expose the surgical site. A small 1 cm incision was conducted on the dorsal neck, and a subcutaneous tunnel to the flank was created. The Ang II-containing osmotic pump was pushed into the flank through the subcutaneous tunnel. The incision was closed using interrupted sutures. Mice were administered subcutaneous buprenorphine (0.05mg/kg), twice per day, for pain control during the first three days and then as needed. Mice continued on the WD for an additional 4 weeks.

Vascular ultrasonography was performed on the first set of mice (14 WT and 13 *Tymp^−/−^*) to monitor the dynamic changes of the AA. We monitored the changes in the aortic inner diameter, blood flow, and other special findings such as thrombus formation, hematoma formation, obvious wall thickness, and calcification, before, 2 weeks, and 4 weeks after the minipump implantation. The inner diameter of the AA was measured at the suprarenal level. Flow parameters, including peek and mean velocity, time range of pulse, as well as acceleration and deceleration of pulse, which are affected by the aorta size or the presence of hematoma, were collected.

### 2.3. Gross examination of AAA

Four weeks after chronic Ang II perfusion, mice were anesthetized with isoflurane, and laparotomy was carried out through the middle abdominal incision. Euthanasia was carried out by drawing whole blood from the inferior vena cava. Subsequently, mice were perfused with 10 mL of 10% formalin through the left ventricle to remove residual blood from the vessels. Abdominal organs and adipose tissues were then removed. If an aneurysm was confirmed visually, the soft tissues surrounding the aorta were cleaned, and a photograph of the AAA was taken. The entire aorta, including major branches, was isolated by careful dissection from surrounding tissues and fixed in 10% formalin for 48 hours. The AA with an aneurysm was sectioned into three segments and embedded in paraffin. In a separate group of mice (12 WT and 13 *Tymp^−/−^*), perfusion with cold PBS was performed, and aorta segments were directly embedded in OCT for frozen sectioning. Sections of six micrometers were cut and utilized for histological examination or in situ zymography.

### 2.4. Histology examination

Hematoxylin and eosin (H&E), Elastica van Gieson (EVG) staining, and Trichrome staining were performed to assess vascular structure, elastin integrity, and fibrosis. Standard immunohistochemical (IHC) or immunofluorescent staining was conducted using antibodies as detailed in the Results section.

As listed in Supplementary Table 1, a total of sixteen AAA samples were utilized for histological examination: eight obtained from Chinese patients (#1 to #8) at the Guangdong Cardiovascular Institute and eight obtained from Caucasian patients (#9 to #16) at the University of Missouri School of Medicine. These samples were analyzed for the expression of TYMP, α-smooth muscle actin (α-SMA), phosphorylated AKT (p-AKT), and transforming growth factor beta 1 (TGFβ1). Image scoring was performed based on representative images presented in **Supplementary Figure 1**.

The human studies were approved by the Institute Research Committee of Guangdong General Hospital (IRB#: GDREC2016255H, PI: Qiuxiong Lin) and by the Institutional Review Board of the University of Missouri School of Medicine (IRB#2026026, PI: Shiyou Chen). Written informed consent was obtained from all subjects.

### 2.5. qPCR

Total RNA was isolated from the human aortic vessel wall, frozen AAA samples (AAA #1 to #8) and healthy donors, or VSMCs using the Qiagen Universal RNA extraction kit. One microgram of total RNA was used for cDNA construction using the Super Script VILO cDNA Synthesis Kit (Thermofisher Scientific, Waltham, MA). **Supplemental Table 2** lists the primers used for analyzing the genes targeted. The PowerUp™ SYBR™ Green Master Mix (ThermoFisher) kit was used for qPCR analysis using the ABI SimpliAmp Thermal Cycler.

### 2.6. Cell culture

A rat VSMC cell line, C2, stably expressing human TYMP, and the control cell line, PC, were established in previous studies^10, 12^. Additionally, VSMCs were cultured from WT and *Tymp^−/−^* mice aorta using an explant method^10^, and cells from passages 4 to 8 were used in this study. All cells were cultured in full culture media (FCM) composed of Dulbecco’s Modified Eagle Medium (DMEM), 10% fetal bovine serum, and antibiotics.

### 2.7. To study the impact of platelets on VSMC proliferation

Platelets were isolated from WT and *Tymp^−/−^* mice and washed to remove plasma components. Serum-starved WT VSMCs were stimulated with FCM in the presence of WT or *Tymp^−/−^* platelets at a density of 10^7^ platelets/well for 12 and 24 hours, and then cell proliferation was assessed using an MTT assay.

### 2.8. Impact of TYMP on MMP expression and activation in VSMCs

C2 and PC cells were cultured in FCM for 8 hours, washed with PBS, and then incubated in serum-free DMEM for 18 hours. The media were collected for gelatin zymography^13, 16^. Subsequently, the serum-starved cells were treated with serum-free DMEM containing TNF-α (10 ng/ml) for various durations. Cells were lysed in RIPA buffer containing protease and phosphatase inhibitors for western blot assay. In another set of experiments, serum-starved PC and C2 cells were treated with serum-free DMEM in the presence or absence of 1 μM Ang II for 24 hours. The culture media were collected for zymography, and the cells were lysed in RIPA buffer for western blot assays. Additionally, serum-starved C2 and PC cells were treated with the thrombospondin-1 (TSP1) type 1 repeat domain (TSR)^17^ or tipiracil, a selective TYMP inhibitor, for different durations. Conditioned media were collected for zymography, and cells were collected for western blot assays.

### 2.9. In situ zymography

The AA embedded in OCT were sectioned and mounted on a slide glass. The sections were washed with PBS and then incubated with a reaction buffer containing 25 µg/mL fluorescein-conjugated DQ gelatin (D12054, ThermoFisher Scientific) for 18 hours at room temperature in a dark, humid slide incubation box^18^. Images were captured under conditions that eliminated background autofluorescence, utilizing *Tymp^−/−^* sections incubated with reaction buffer only. The mean fluorescent intensity of each 20x image (green channel only), as well as the mean background intensity (without any tissues), was analyzed with ImageJ. The data were presented as the whole image mean minus the background mean.

### 2.10. Mouse plasma cytokine array

Mouse plasma was pooled from randomly selected 6 WT mice (including 3 mice with confirmed AAA) or 6 *Tymp^−/−^* mice (including one with AAA and 5 randomly selected). This pooled plasma were then used for determining plasma cytokine levels using the Proteome Profiler Mouse Cytokine Array Kit, panel A. The membrane images were scanned and analyzed using ImageJ.

### 2.11. Statistics

The data were analyzed using GraphPad Prism (version 10.1.2) and expressed as Mean ± SE. Data normality and equal variance were assessed using the D’Agostino-Pearson normality test to justify the use of the 2-tailed Student’s *t-test*, Mann-Whitney test, or One- or two-way ANOVA for comparisons of two or more groups or factors. Mixed effects two-factor ANOVA was utilized for multiple time points and two-factor comparisons. For the mixed effects model, we treated genotype as the random effect and time as the fixed effect. A *p* ≤ *0.05* was considered statistically significant.

## 3. Results

### 3.1. TYMP expression is increased in the AAA vessel wall of patients

To assess the involvement of TYMP in AAA pathogenesis, we studied 16 AAA patients and presented their clinical data in **Supplementary Table 1**. Compared to the aortic walls of healthy donors, AAA patient samples showed significant structural disruption, as demonstrated by H&E and Masson’s Trichrome stains (**Fig. 1A and 1B**). We observed a disorganized VSMC layer, with the accumulation of blood cells, plaque, fatty deposits, and connective tissue. We also observed a pronounced increase in disoriented fibrotic tissue, compromising the structural integrity of the VSMC layer.

**Fig. 1.**
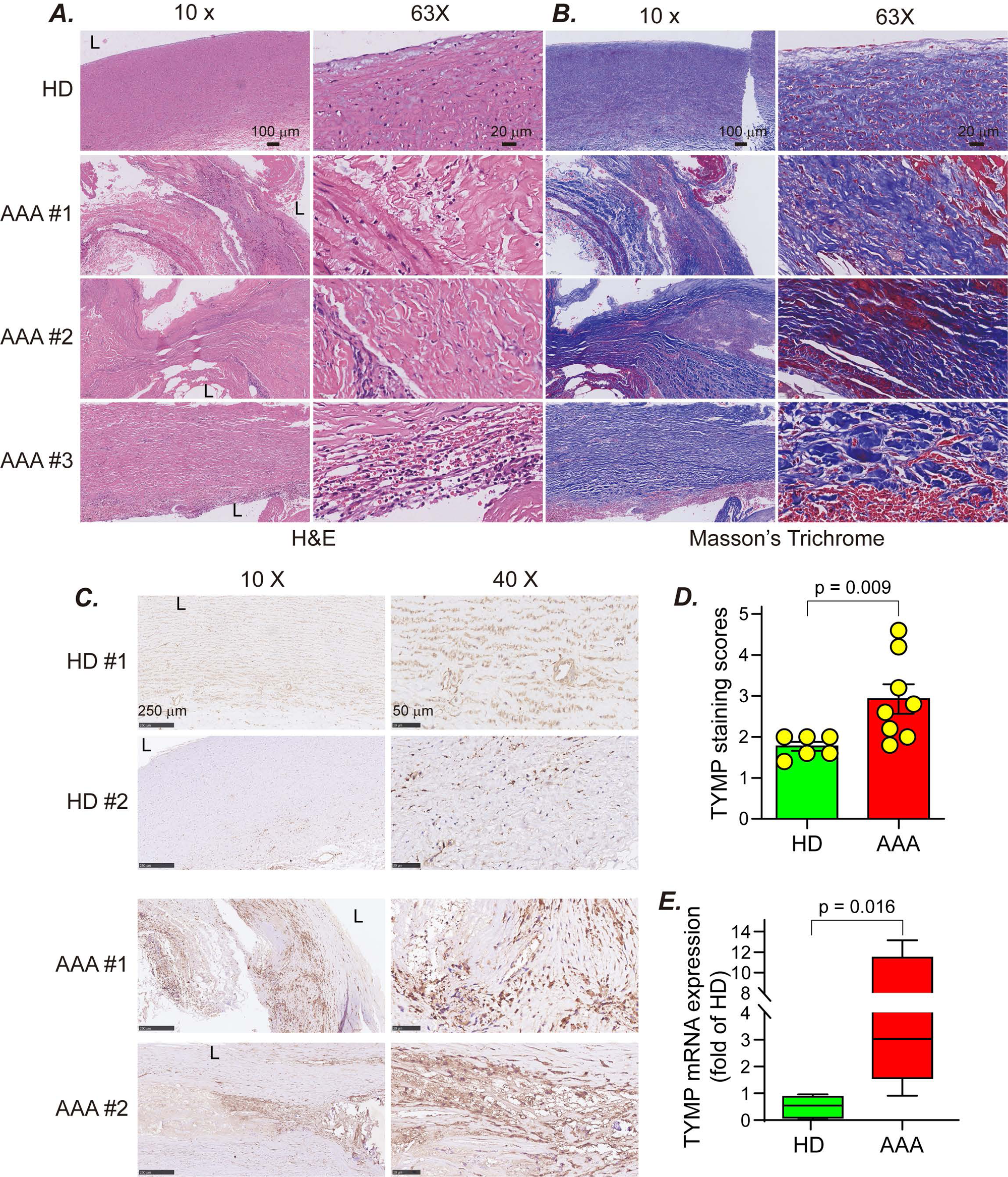
TYMP is increased in the human abdominal aortic aneurysm vessel wall. Human AAA vessel wall and normal abdominal aorta from healthy donors (HD) were sectioned and stained with ***A***, H&E, ***B***, Masson’s Trichrome staining, and ***C,*** IHC for TYMP. Brown in C indicates a positive stain. ***D.*** TYMP staining scores. ***E.*** qPCR analysis of TYMP expression in the aortic vessel wall harvested from HD and AAA patients. N = 6, in health donors and 8 in AAA.

Consequently, we compared TYMP expression in normal aorta and AAA by IHC and qPCR. As shown in **Fig. 1C**, TYMP expression was detectable in both healthy aortas and AAA vessel walls; however, the staining intensity and score were significantly higher in the aneurysm samples (**Fig. 1D**). The majority of the TYMP was expressed within the vasa vasorum in the healthy aorta. However, TYMP was expressed by all cells in the AAA vessel wall, with a significant increase observed in VSMCs (**Supplementary Figure 2**). qPCR further confirmed the increase of TYMP mRNA in AAA samples when compared to the healthy aortas (**Fig. 1E**). These data suggest that TYMP expression is increased in the AAA vessel wall compared to healthy controls.

### 3.2. TYMP deficiency in mice reduces the prevalence of AAA

To investigate the role of TYMP in AAA development, we chronically perfused Ang II into WD-fed WT and *Tymp^−/−^* mice, composing 26 animals in each genotype. Five WT and two *Tymp^−/−^* mice died prematurely, with the majority of these deaths occurring within 10 days after Ang II perfusion. As illustrated in **Fig. 2A**, the *Tymp*^−/−^ group exhibited a lower, albeit statistically non-significant, mortality rate. Autopsies revealed hemorrhage in the thoracic cavity (**Supplementary Figure 3**). Heart rupture was not confirmed, suggesting an aortic rupture; however, due to advanced autolysis and decomposition of the carcasses, the bleeding site could not be determined. These mice were excluded from the subsequent analysis of AAA prevalence.

**Fig. 2.**
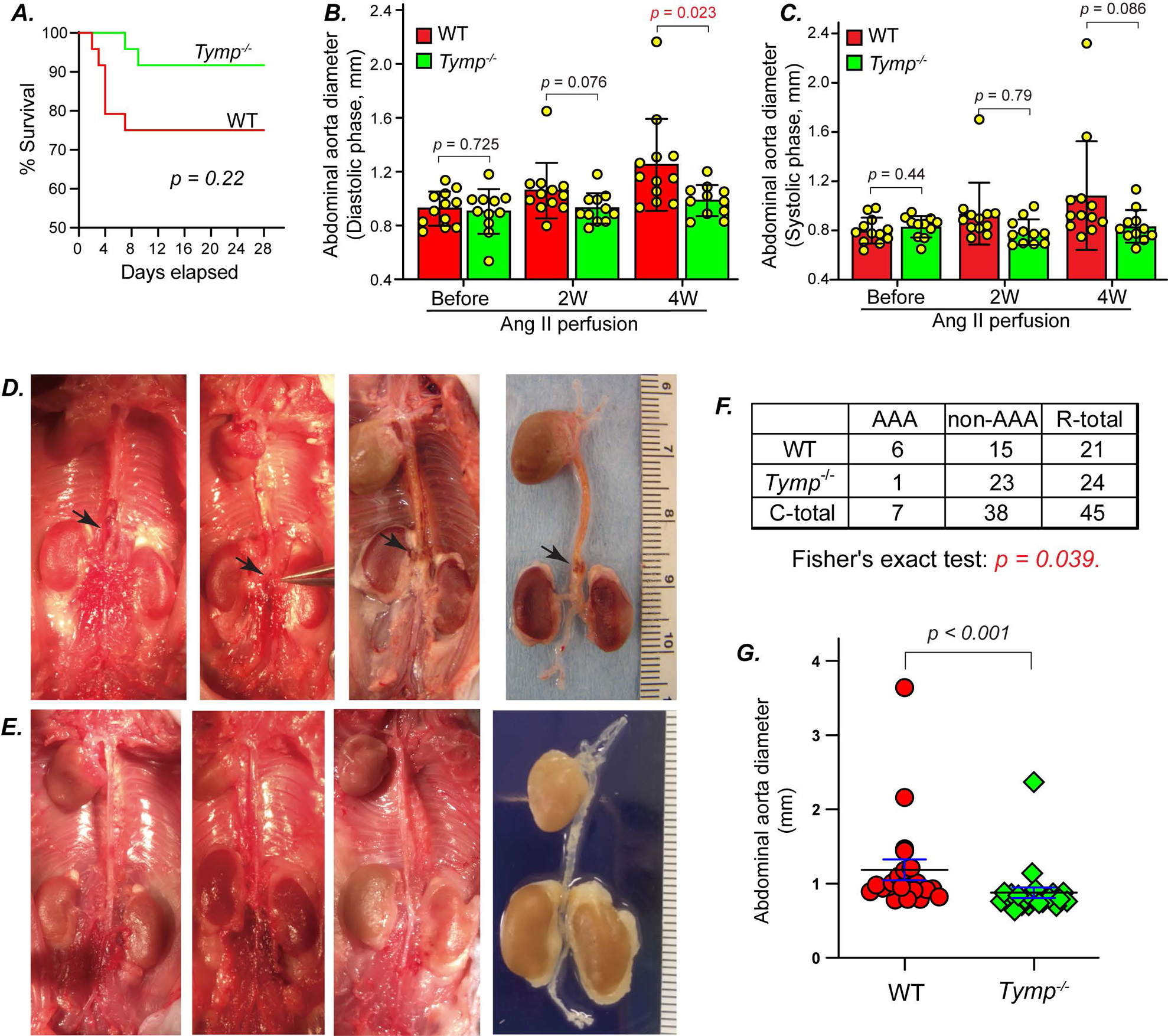
TYMP deficiency reduced the prevalence of AAA in mice. WT and *Tymp^−/−^* mice were fed a Western diet starting at 4 weeks of age for 8 weeks, and then received chronic Ang II perfusion in a dose of 1 µg/kg/day delivered with an Alzet® osmotic mini pump for 4 weeks. The development of AAA was monitored with echography. ***A.*** Survival curve analysis, n = 26. ***B.*** Inner diameter of abdominal aorta (AA) in the diastolic phase. ***C.*** Inner diameter of AA in the systolic phase. ***D.*** Gross finding of aortic tree in the WT mice. Arrows indicate AAA. ***E.*** Gross finding of aortic tree in the *Tymp^−/−^* mice. ***F.*** Contingency table showing mice number based on whether AAA formation was observed by necropsy. Fisher’s exact test was used for statistical analysis. ***G.*** AA diameter at the suprarenal levels measured by caliper, n=21 in WT and 24 in *Tymp^−/−^* group.

As shown in **Fig. 2B** and **Supplementary Figure 4A** and **4C**, perfusion of Ang II for 4 weeks significantly increased the luminal diameter of the AA during the diastolic phase in WT mice, rising from an average of 0.925 ± 0.127 mm before Ang II administration to 1.25 ± 0.342 mm post-treatment (*p* = 0.006, n = 12). However, this effect was not observed in *Tymp^−/−^* mice, where the AA luminal diameter remained similar before and after Ang II infusion (0.904 ± 0.166 mm pre-Ang II vs. 0.984 ± 0.116 mm at 4 weeks, *p* = 0.2). Consequently, the post-treatment diastolic luminal diameter in *Tymp*^−/−^ mice was significantly smaller compared to that in WT mice (**Fig. 2B**). Additionally, while not reaching statistically significant, there was an observed trend towards increased AA luminal diameter in the systolic phase for WT mice, as depicted in **Fig. 2C** and Supplementary Figure 4B and **4D**, a trend not seen in *Tymp*^−/−^ mice.

Necropsy examination confirmed the presence of AAA in 6 out of 21 WT mice, resulting in an incidence rate of 28.6%, while in the *Tymp^−/−^* cohort, only 1 of 24 mice developed an AAA, representing a 4.2% incidence (**Fig. 2D and E**). Fisher’s exact test indicated that TYMP deficiency significantly reduced the prevalence of AAAs (**Fig. 2F**, *p* = 0.039). The identified aneurysms exhibited a fusiform shape, typically located at the suprarenal artery, with their longitudinal axis aligning with the artery. The average AA diameter measured above the renal artery was larger in WT mice compared to *Tymp*^−/−^ mice (**Fig. 2G**). The aneurysms appeared dark red, indicative of intramural thrombosis. It is important to note that although sudden deaths were attributed to hemorrhage in the thoracic cavity, no thoracic aortic aneurysms were confirmed in any surviving mice.

In addition, we evaluated the hemodynamics in the AA, including peak and mean velocities, as well as acceleration and deceleration patterns of blood flow before-, and 2-, and 4-week post-Ang II infusion. Data from **Supplementary Figure 5A** to **5D** suggest that TYMP deficiency had no significant impact on these flow parameters at the evaluated time points. In contrast, among the three WT mice with confirmed AAAs, there was a significant increase in acceleration at 2- and 4-week post-infusion, as displayed in Supplementary Figure 5E (WT panel) and **5F**.

### 3.3. TYMP deficiency attenuates the distortion of the aortic wall in the murine AAA model

H&E staining of WT mouse AAA cross-section demonstrated dilated lumens (**Supplementary Figure 6A**). Within these aneurysms, we observed either freshly formed (WT#1) or fibrotic (WT#2) hematoma, which disrupted the media and adventitia of the affected aorta. In contrast, the aortic structure of *Tymp^−/−^*mice remained unchanged, preserving wall integrity even in the mouse with an aneurysm. Subsequent EVG staining highlighted that the elastic fibers within the vessel walls of *Tymp^−/−^* mice retained better integrity, contrasting with the breakage and disarray seen in the elastic fibers of the aortic wall of WT mice (**Supplementary Figure 6B**).

Double immunofluorescence staining for vWF, expressed in both endothelial cells and platelets (and thus platelet-rich thrombus), and α-SMA, the VSMC marker, showed disruption of endothelial layer in the aortas with aneurysm (**Fig. 3A**). The high amount of vWF in aneurysmal hematoma areas indicated platelet-rich thrombus formation. The aorta of *Tymp*^−/−^ mice maintained a normal structure across all layers, consistent with H&E and EVG staining results. The α-SMA positive signal found within the hematoma was verified through IHC (**Fig. 3B**), and Masson Trichrome staining showed these regions as fibrotic (in blue) interwoven with blood cells (in red) (**Fig. 3C**). This suggests that the α-SMA positive cells within the hematoma may be myofibroblasts involved in the pathological response to aneurysm formation or healing. No CD68-positive macrophage accumulation was observed in both WT and *Tymp^−/−^* vessels (Data not shown).

**Fig. 3.**
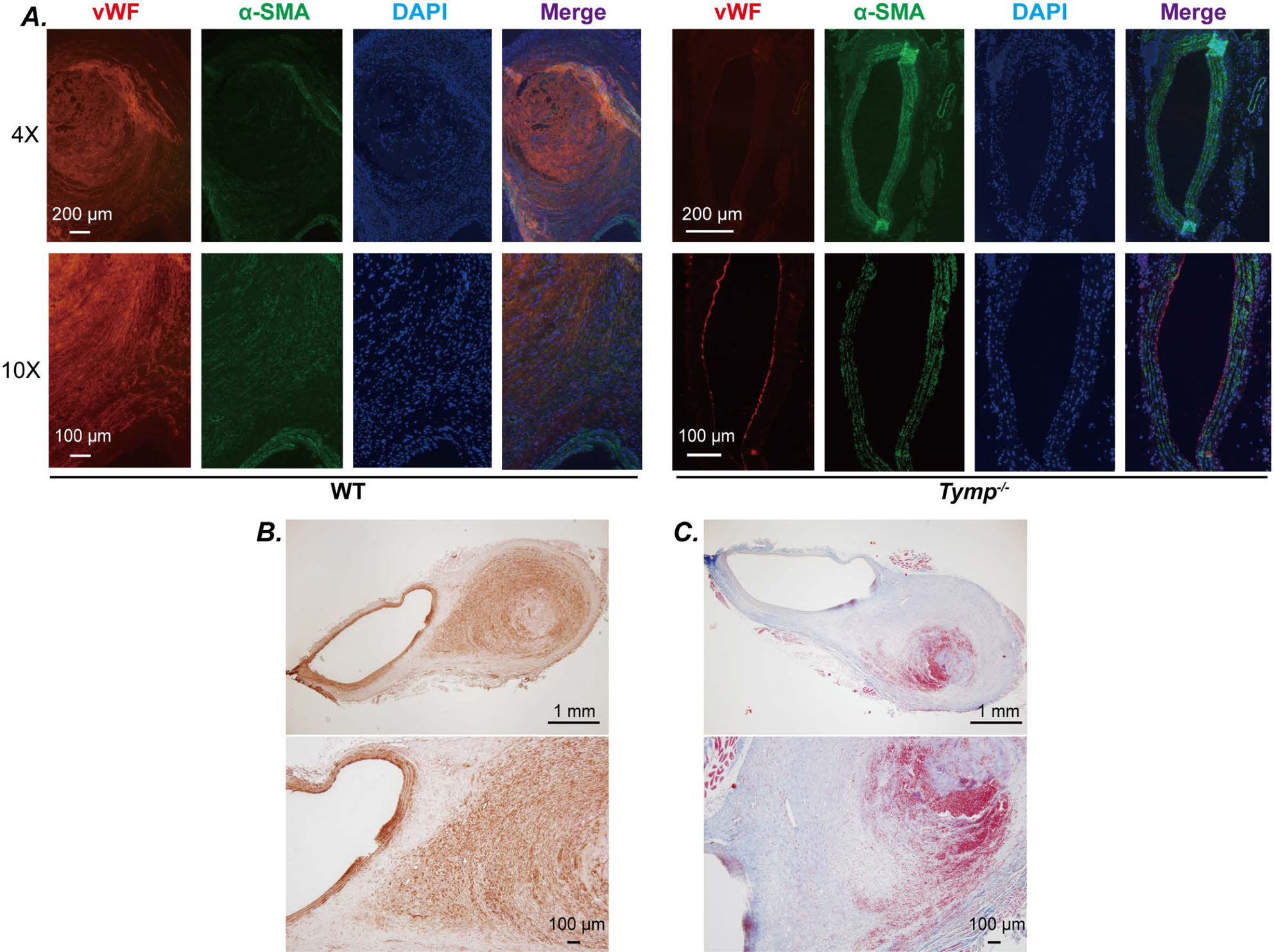
TYMP deficiency attenuates vessel wall structural distortion in the murine AAA model. ***A.*** Paraffin-embedded AAA vessel walls from WT mice and AA vessel walls from *Tymp^−/−^* mice were sectioned and double immunofluorescence staining for vWF and a-SMA was conducted. Nuclei were stained with DAPI. ***B.*** IHC of α-SMA in the AAA section. Brown indicates positive staining. ***C.*** Masson’s trichrome staining of the AAA section.

### 3.4. TYMP enhances MMP2 production and secretion in VSMCs

MMPs, particularly MMP2 and MMP9, are known to play crucial roles in AAA formation^6, 19^. TYMP expression has been shown to correlate positively with MMP2 and MMP9^13, 20^. Consequently, we measured MMP2/9 activity in C2- and PC-cell-conditioned media by gelatin zymography and found that overexpressing TYMP significantly increased MMP2 activity (**Fig. 4A**). MMP9 was undetectable under these conditions. However, intracellular levels of MMP2 were substantially reduced in C2 cells when assessed via western blot analysis (**Fig. 4B**). We thus examined the expression of MMP2 at the mRNA levels and found it was significantly increased in C2 cells (**Fig. 4C**), aligning with its changes in activity. These data indicate that TYMP promotes both MMP2 transcription and secretion in VSMCs.

**Fig. 4.**
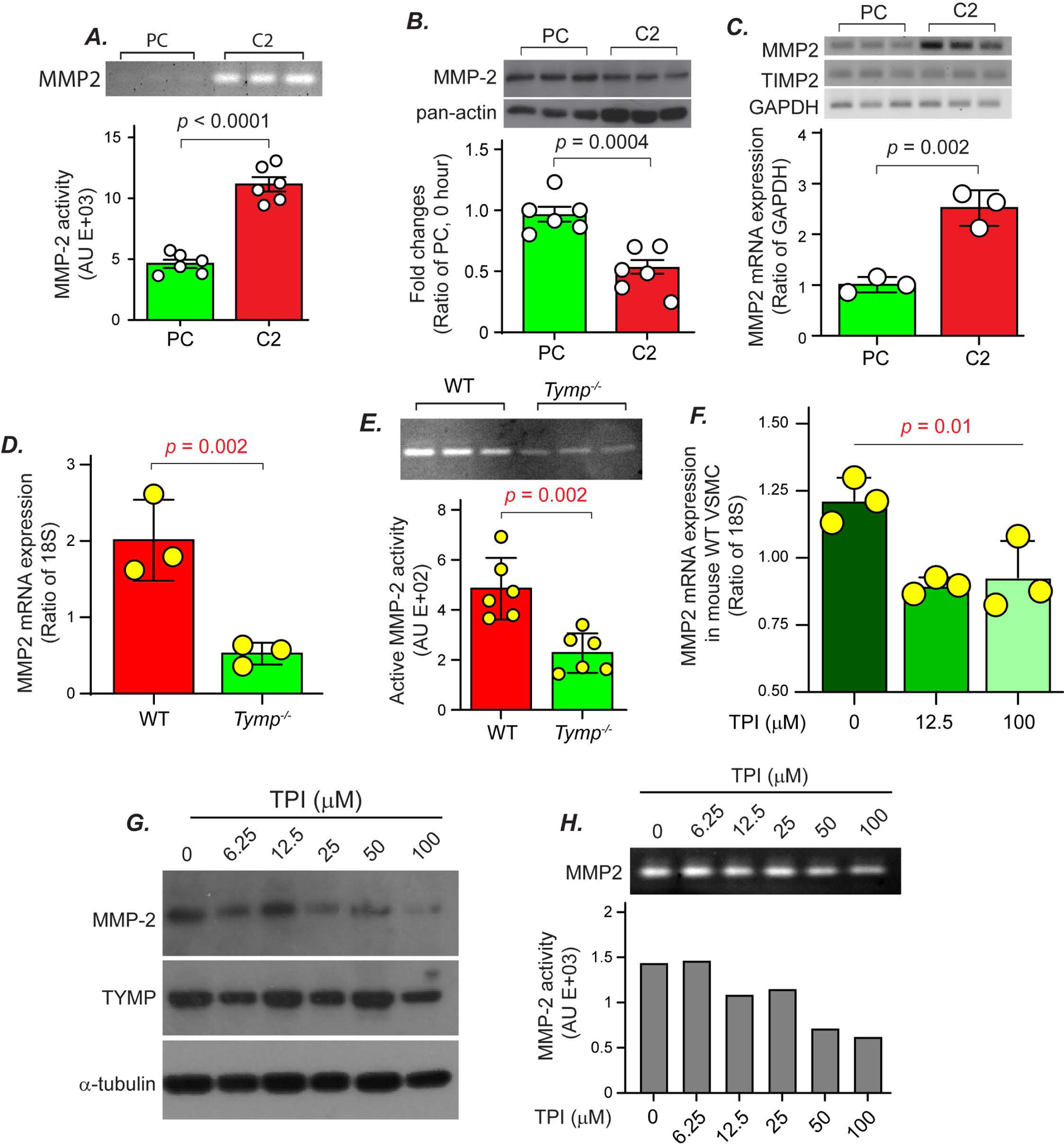
TYMP enhances MMP2 production and secretion in VSMCs. C2 and PC cells were cultured in a 6-cm plate for 8 hours, washed with warm PBS, and then incubated in serum-free DMEM for 24 hours. The media were collected and used for a gelatin zymography assay (***A***). Cells were harvested for Western blot assay (***B***) or RT-PCR (***C***). ***D.*** WT and *Tymp^−/−^* VSMCs were cultured in normal conditions, and total RNAs were extracted and used for qPCR assay. ***E.*** WT and *Tymp^−/−^*VSMCs were seeded into 6-well plates and cultured for 24 h. The cells were rinsed with PBS and then cultured in serum-free DMEM for another 24 h and then media were collected for Zymography assay. ***F.*** WT VSMCs were seeded into 6-well plates and cultured overnight. The cells were rinsed with PBS and then cultured in serum-free media in the presence or absence of TPI for another 24 h. Cells were collected for qPCR assay of MMP2 expression. ***G*** & ***H***. C2 cells were cultured in serum-free media in the presence of different concentrations of TPI for 24 h, and then cells and media were collected for western blot (***G***) and zymography (***H***) assay of MMP2 expression.

These results were corroborated using VSMCs primarily cultured from WT and *Tymp*^−/−^ mice. As shown in **Fig. 4D and E**, TYMP deficiency markedly decreased MMP2 mRNA expression and its activity in these cells. Tipiracil treatment also dose-dependently reduced MMP2 expression at mRNA, protein, and activity levels (**Fig. 4F, 4G, and 4H)**. Taken together, these data suggest that TYMP plays an important role in enhancing MMP2 expression and secretion, which could contribute to the development and progression of AAA.

### 3.5. TYMP enhances the expression of pro-inflammatory cytokines

Pro-inflammatory cytokines and chemokines, including tumor necrosis factor-α (TNF-α)^21^, interleukin (IL)-1β^22^, IL-17^23^, IL-23^24^, etc., have been implicated in AAA formation and disease progression. We recently demonstrated that TYMP expression is increased in COVID-19 patients and its expression is significantly correlated with COVID-19-associated inflammation and thrombosis^25^. To determine TYMP’s role in promoting inflammation and subsequent AAA development, we performed a Proteome Profiler Mouse Cytokine Array on plasma pooled from six WT and six *Tymp*^−/−^ mice. The analysis, as depicted in **Fig. 5A** and **Supplementary Figure 7**, revealed elevated levels of various cytokines and chemokines in WT plasma, many of which are established contributors to AAA formation. Interestingly, the levels of IL-1α and the tissue inhibitor of metalloproteinases 1 (TIMP1) were found to be significantly lower in the WT mice plasma. The suppression or genetic elimination of IL-1α or TIMP1 has been associated with the progression of AAA^26, 27^.

**Fig. 5.**
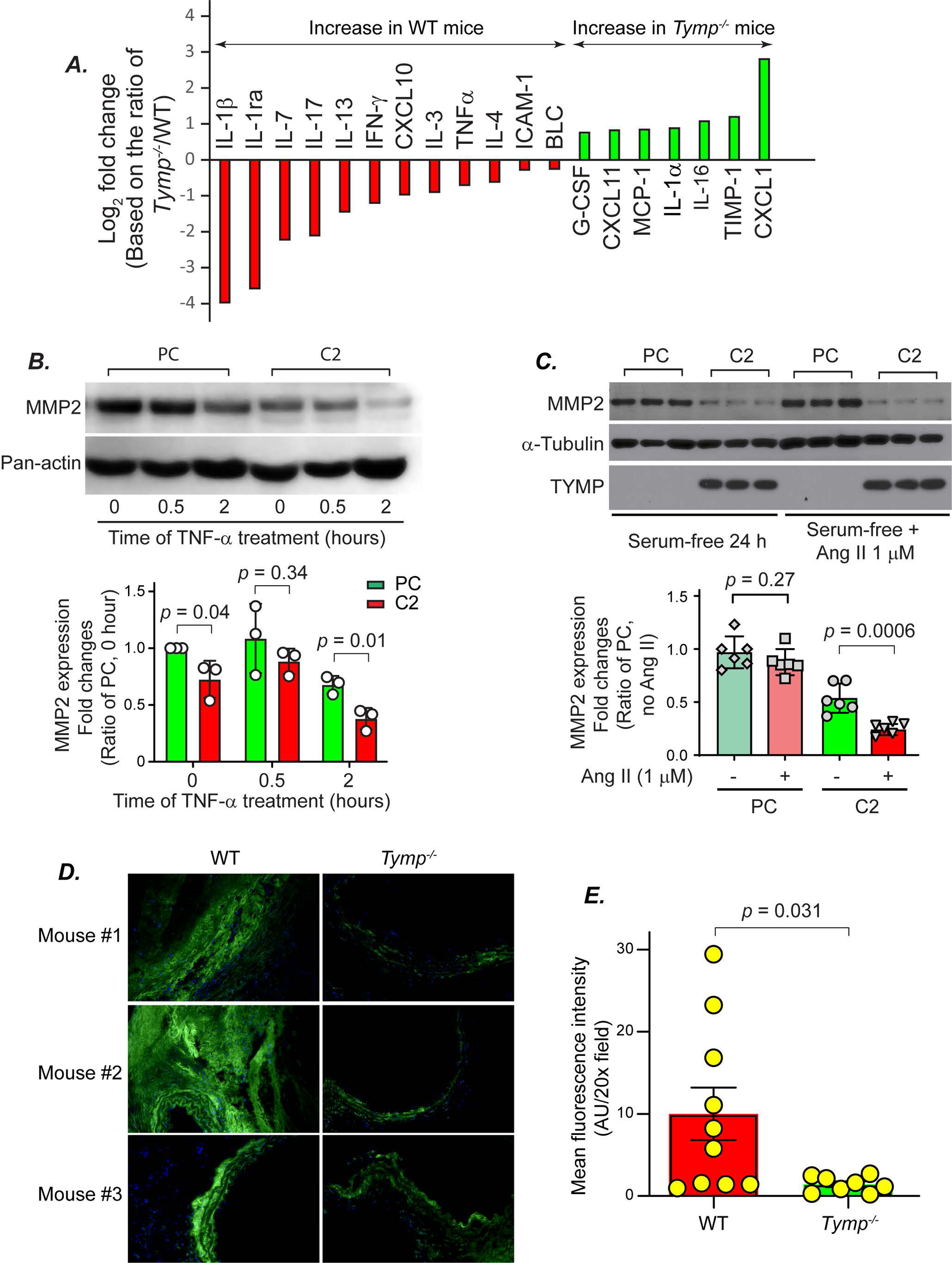
TYMP is pro-inflammatory and enhances MMP2 production and secretion in VSMCs. ***A.*** Plasma pooled from 6 WT mice (including 3 mice with AAA and 3 randomly selected) and 6 *Tymp^−/−^* mice (including 1 mouse with AAA and 5 randomly selected) were subjected to the Proteome Profiler Mouse Cytokine Array, Panel A. The intensity of each spot was analyzed by ImageJ. The ratio of *Tymp^−/−^* to WT was calculated, and the average value of each protein was presented as Log2 fold changes. Molecules with changes of more than 25% were shown in the graph. ***B.*** Serum-starved C2 and PC were treated with TNF-a (10 ng/mL) in serum-free media for the indicated times and then cells were harvested for western blot assay. ***C.*** PC and C2 cells were serum-starved for 24 hours and then treated with 1 mM Ang II in serum-free DMEM for 24 hours. Cells were harvested for western blot assay. ***D & E***. Three AAA from the WT mice, as well as one AAA (#3) and two randomly selected AA (#1 and 2) from the *Tymp^−/−^* mice were embedded in OCT, sectioned into 6 µm slices, and mounted onto a slide glass. The AA tissues were washed with PBS and then incubated with fluorescein conjugated DQ gelatin for 18 hours. Representative images are shown in *D*. The mean fluorescence of the image, which represents the activity of MMPs in the vessel wall, was analyzed by ImageJ and used for statistical analysis (*E*).

To investigate the relationship among TYMP, inflammation, and MMPs, we treated serum-deprived PC and C2 cells with TNF-α and assessed MMP2/9 expression and activation. As shown in **Fig. 5B**, baseline intracellular MMP2 levels were naturally lower in C2 cells. Following TNF-α exposure for 30 minutes, a slight uptick in MMP2 expression was observed, followed by a substantial decrease at the 2-hour mark in both cell types, indicating increased MMP2 secretion. Two hours post TNF-α treatment, MMP2 activity became detectable in C2 cells (albeit undetectable in PC cells), as evidenced by a very faint signal on zymography (**Supplementary Figure 8A** with a yellow arrow), This implies that TNF-α stimulates MMP2 secretion, which is further augmented by TYMP.

Additionally, to probe the combined effects of TYMP and Ang II on MMP2 expression, we administered 1 µM Ang II to serum-free PC and C2 cells over 24 hours. According to **Fig. 5C**, Ang II had no significant impact on MMP2 levels within PC cells. However, in C2 cells, Ang II treatment moderately but significantly reduced intracellular MMP2 levels, suggesting an increase in MMP2 secretion. This effect, however, was not verified by zymography (**Supplementary Figure 8B**).

In vivo examination through in situ zymography of aortic sections from WT mice with confirmed AAAs and from *Tymp*^−/−^ mice, specifically one with AAA and two without, corroborated the in vitro findings. TYMP deficiency was shown to substantially decrease MMP activity in the aortic walls, as displayed in **Fig. 5D and E**, further supporting the association between TYMP, inflammation, and AAA development.

### 3.6. TYMP enhances AKT phosphorylation in VSMCs

The presence of intraluminal thrombus is a regular occurrence in AAAs, but its effect on AAA enlargement remains incompletely understood. Platelets are known to contain high levels of TYMP^9, 28^. To investigate the direct impact of platelets on VSMC activity, we co-cultured WT VSMCs with both WT and *Tymp^−/−^* platelets, which were washed to remove plasma components. Consistent with previous findings, WT platelets appeared to suppress VSMC proliferation, as indicated in **Fig. 6A**. These data suggest that TYMP, whether originating from platelets or other cellular sources, can impact VSMC functionality and potentially contribute to vessel wall weakening in certain pathological conditions.

**Fig. 6.**
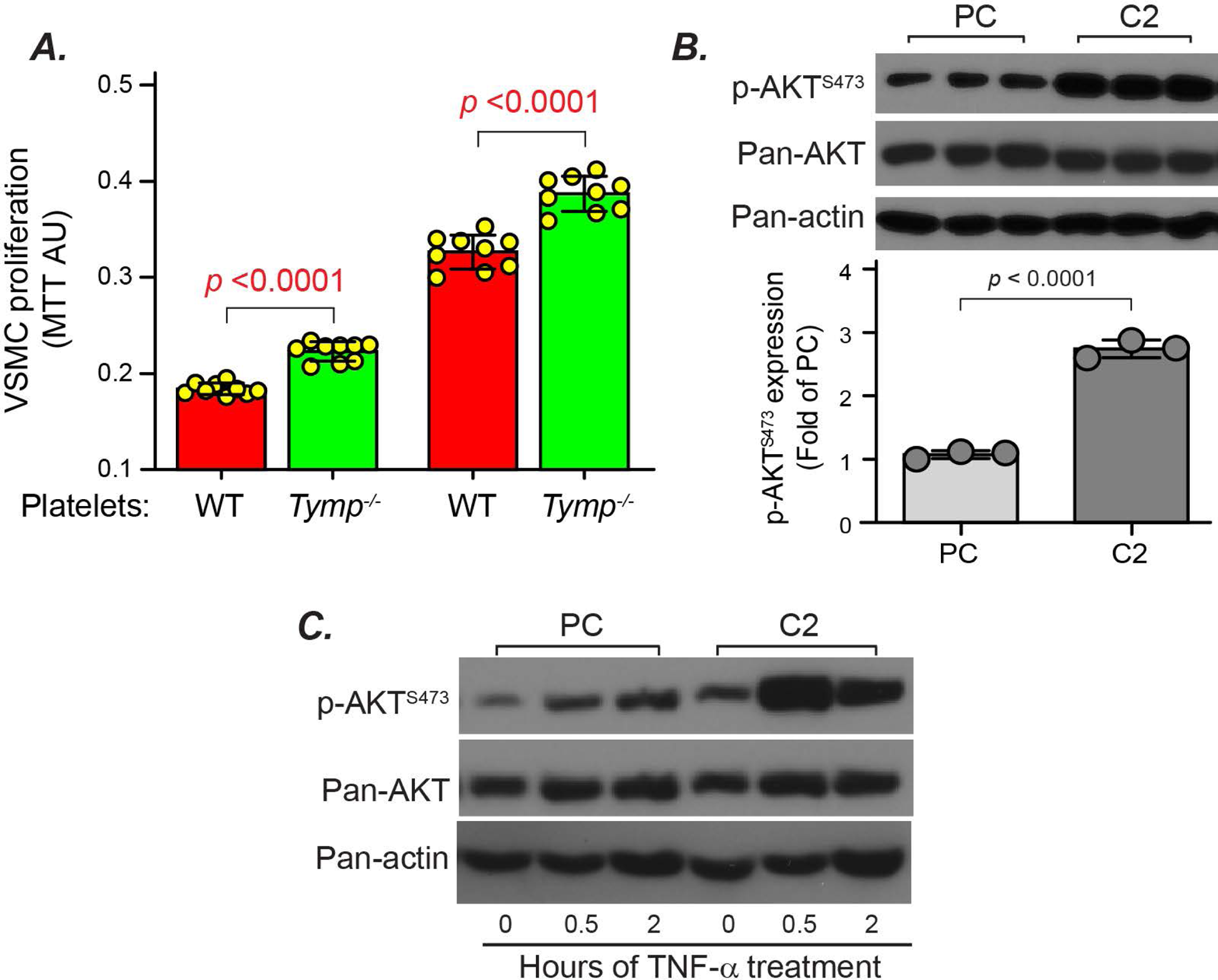
TYMP-expressing platelets inhibit VSMC proliferation and TYMP overexpression leads to constitutive AKT activation in VSMCs. ***A.*** Murine WT VSMCs cultured in a 96-well plate were serum-starved overnight and then stimulated with FCM containing WT or *Tymp^−/−^* platelets (10LJ platelets/well). Cell proliferation was assessed 12 and 24 hours later using an MTT assay. ***B.*** C2 and PC cells were cultured in FCM overnight and then cells were harvested for Western blot assay of AKT activation and expression. ***C.*** Serum-starved C2 and PC cells were treated with TNF-α for the indicated durations and cell lysates were used for western blot assay of AKT activation. Note: The Pan-actin loading control is the same as Fig. 5A.

The mechanism by which TYMP modulates MMP2 expression remains unclear. Our previous research involving platelets and mesenchymal stem cells hinted at a multifaceted interaction between TYMP and the AKT signaling pathway^8, 9^. We found that ADP-stimulated AKT phosphorylation was slightly higher in the early phase but was dramatically decreased in the late phase activation of *Tymp^−/−^* platelets^8^. Conversely, TYMP-deficient mesenchymal stem cells exhibited continuous AKT phosphorylation, even though MMP2 activity was reduced in these cells^20^. These findings imply a dual role for TYMP in AKT pathway regulation: it may generally inhibit AKT activity, but under certain conditions, it enhances AKT activation via an alternate pathway. As shown in **Fig. 6B**, we found that overexpression of TYMP in VSMCs significantly increased AKT phosphorylation at S473. TNF-α stimulated AKT activation in both cell types, with a more pronounced effect in C2 cells (**Fig. 6C** and **Supplementary Figure 9)**. Given the role of AKT activation in promoting MMP2 production in VSMCs and its significance in AAA development^29^, these data suggest that TYMP-enhanced MMP expression could be mediated by the AKT pathway. This hypothesis is further supported by examining AKT activation in human AAA samples, which showed strong p-AKT staining, indicating an increase in AKT activity compared to healthy controls (**Supplementary Figure 10**).

### 3.7. TYMP enhances TGF**β**1 signaling activation and connective tissue growth factor (CTGF) expression in VSMCs

MMP2 activates TGFβ1 signaling and plays an important role in arterial aging, a risk factor for developing AAA. TGFβ1 expression has been significantly increased in the plasma of AAA patients^30^ or patients with aortic dilatation^31^. However, the role of TGFβ1 in AAA development in mice is controversial, with studies reporting both beneficial^32^ and detrimental^33^ effects. To elucidate the involvement of TGFβ1 in AAA pathogenesis, we analyzed its expression in human AAA tissues. As demonstrated in **Fig. 7A, B**, and **Supplementary Figures 11** and **12,** our results revealed a significant upregulation of TGFβ1 within the AAA vessel walls. Notably, TGFβ1 levels exhibited a positive association with TYMP expression (**Fig. 7C**), particularly in VSMCs (**Supplementary Figure 12**).

**Fig. 7.**
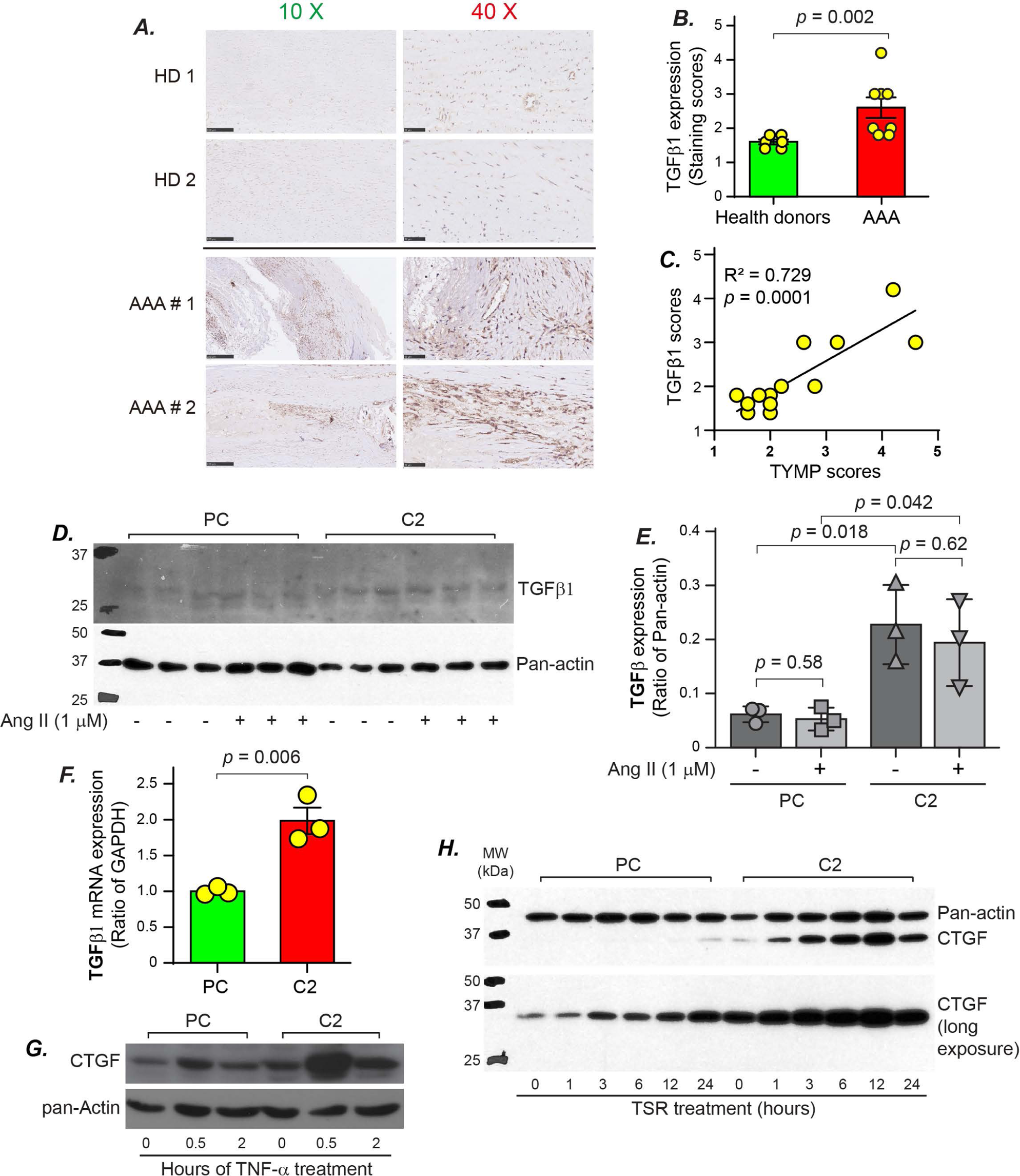
TYMP increases TGFβ1 expression in human AAA and enhances TGFβ1 signaling activity in VSMCs. ***A.*** IHC of TGFb1 in human AAA. ***B.*** TGFb1 staining scores in human AAA. ***C.*** Correlation analysis between TYMP and TGFb1 in human AAA. ***D.*** Serum-starved PC and C2 cells were treated with serum-free DMEM with/without 1 mM Ang II for 24 hours and TGFb1 expression was examined by western blot. ***E.*** Statistical analysis of western blot band intensity showed in D. ***F.*** TGFb1 mRNA expression in PC and C2 cells under normal culture. ***G***. Cell lysates mentioned in Fig. 6C were used for the examination of CTGF expression. ***H.*** Serum-starved PC and C2 cells were treated with serum-free media in the presence of 10 µg/mL TSR for the indicated durations. CTGF was examined by Western blot. Pan-actin was blotted in the same membrane without stripping the CTGF signal as loading control.

To further investigate the regulatory role of TYMP on TGFβ1 expression in VSMCs, we assessed its expression under both baseline and Ang II-stimulated conditions. While Ang II had no discernible impact on TGFβ1 levels in both PC and C2 cells, a marked increase in expression was observed in C2 cells (**Fig. 7D and E**). This elevation in TGFβ1 expression in C2 cells was further validated by qPCR analysis (**Fig. 7F**). These findings suggest that TYMP may play a crucial role in enhancing TGFβ1 expression in VSMCs, thereby potentially contributing to AAA development.

CTGF is a key mediator of tissue remodeling and fibrosis through its interactions with TGFβ1 and integrin αvβ3. The upregulation of CTGF is associated with the pathogenesis and progression of ascending thoracic aortic aneurysm^34^. Activation of integrin αvβ3 plays an important role in the TYMP-mediated proangiogenic process^35^. We thus investigated whether CTGF mediates TYMP’s effect in response to stimulation of TNF-α and thrombospondin-1 (TSR), which activates the TGFβ1 signaling pathway. As depicted in Fig. 7G, TNF-α induced a greater increase in CTGF expression in C2 cells compared to that in PC cells. Similarly, TSR stimulated CTGF expression in a time-dependent manner in both cell lines, with C2 cells showing a significantly higher response (**Fig. 7H**). This evidence supports the premise that TYMP is integral to TGFβ1 signaling and influences the elevation of CTGF expression.

## 4. Discussion

In this study, we utilized a chronic Ang II perfusion model in WT and *Tymp^−/−^* mice on a WD and demonstrated that TYMP plays a pivotal role in AAA development. Our findings revealed an increase of TYMP expression within human AAA vessel walls, and its deficiency in mice significantly reduced Ang II-induced aortic dilation and AAA incidence. Beyond the known inhibitory effect of TYMP in VSMCs^10–12^, we identified additional key contributions of TYMP to AAA prevalence. Specifically, TYMP deficiency markedly altered the expression of inflammatory cytokines and chemokines linked to AAA development. Furthermore, TYMP not only boosts MMP2 production but also its secretion. It triggers constant AKT phosphorylation when overexpressed in VSMCs, and engages in the TSP1/TGFβ1 pathway, elevating TGFβ1 and CTGF levels. This study is the first to highlight TYMP’s essential role in creating a conducive environment for AAA development, a major human vascular disease.

Human AAA formation spans years or even decades influenced by a complex interplay of various pathophysiological elements and risk factors, creating a highly intricate scenario. Consequently, the ideal animal model for studying AAA should accurately mimic both the pathophysiological process and the characteristics of human AAA^36^. While extensive research has explored the impact of genetic factors on AAA development by introducing gene modifications into *Apoe^−/−^* or *Ldlr^−/−^*backgrounds, these models fall short of accurately replicating human AAA pathophysiology. This discrepancy arises because *APOE* or *LDLR* deficiencies are rare in humans, and the occurrence of double gene deficiencies is almost non-existent. Although these studies yield valuable insights into aortopathy etiology, employing models with fewer genetic modifications could offer fresh perspectives more aligned with the pathophysiology of human AAA. In our study, after administering chronic Ang II infusion to mice fed WD for four weeks, we observed a 28.6% incidence of AAA in WT mice. This rate surpasses that of a prior study, where only 20% of C57BL6 mice fed a standard laboratory diet developed AAA^37^. The pathological alterations, morphology, and characteristics of AAA in WT mice closely resembled those in human AAA, indicating that this model effectively replicates the pathophysiology of AAA and is suitable for further study.

Our model’s higher AAA incidence indicates that systemic changes associated with the WD promote AAA development. Our previous and ongoing studies in a different project showed that feeding mice with WD dramatically enhanced thrombosis^8^ and increased TYMP expression in platelets and livers (data not shown), respectively. While the WD is well known to cause systemic chronic inflammation^5^, this study is the first to demonstrate that TYMP plays an important role in WD-induced systemic inflammation. This aligns with several studies indicating TYMP’s role in upregulating inflammatory cytokines like IL-8 and CXCL10^7^, and its positive correlation with plasma C-reactive protein levels, a common clinical inflammation marker^25^. These novel findings about TYMP-mediated alterations of inflammatory cytokines highlight TYMP’s involvement in the AAA pathology. However, additional studies are necessary to elaborate on how TYMP influences the expression of these inflammatory cytokines, especially concurrently.

The exact pathophysiology behind aneurysm formation remains unclear. However, it is widely recognized that VSMCs are the predominant cell type involved and play crucial roles in this process. Dysregulated VSMC function, behavior, and antioxidant status have been linked to vascular diseases, including neointimal hyperplasia, atherosclerosis, and AAA^38^. Studies from our laboratory and others have demonstrated that TYMP plays an inhibitory role in regulating VSMC function^10–12^. In this study, we further demonstrated that WT platelets have a stronger inhibitory effect on VSMC proliferation than *Tymp^−/−^* platelets. Recent studies have demonstrated that the accelerated growth of AAA is associated with platelet activation and thrombosis in aneurysmal segments^39^. Thus, we predict that besides TYMP’s contribution to promoting inflammation, its role in augmenting platelet activation and thrombosis, along with its inhibitory impact on VSMC functionality, collectively creates a conducive environment for the development of AAA.

In a canine transmyocardial laser revascularization model, we showed that laser treatment-enhanced angiogenesis is correlated with the increased expression of TYMP, MMP2, MMP9, and urokinase-type plasminogen activator (uPA)^13, 16^, suggesting a potential association between TYMP, MMP2/9, and uPA. The activity of MMPs is strictly controlled at several levels, including transcription, production, and activation, as well as binding to their natural endogenous inhibitors, TIMPs. While TYMP overexpression does not alter TIMP2 levels (**Supplementary Figure 13**), we observed a reduction in TIMP1 levels in WT plasma, as indicated in Fig. 5A. This observation aligns with in situ zymography results, which demonstrated increased MMP activity in the aortic walls of WT mice.

Various aortopathy has been linked to the gain-of-function mutation or dysregulation of the TGFβ1 signaling pathway^40^. As mentioned above, the role of TGFβ1 in the development of AAA is controversial, and both beneficial and detrimental effects have been reported^32, 33^. A recent review further highlighted that both activation and inhibition of TGFβ1 disrupt its vital role in maintaining normal vascular biology, potentially leading to aneurysm formation by triggering both canonical and non-canonical signaling pathways^40^. Interestingly, the AKT pathway, a non-classical pathway downstream of TGFβ1, is implicated in the development of aortic aneurysms^29^. TGFβ1 is secreted in a latent form by cells, binding to latent TGFβ1 binding protein (LTBP) and the TGFβ1 propeptide (also known as latency-associated peptide, LAP). We found that TYMP not only enhances TGFβ1 transcription but also increases the active form of TGFβ1 in VSMC, confirmed by an increase of the 25 kDa TGFβ1. These in vitro studies reinforce the observed positive correlation between TYMP and TGFβ1 expression in human AAA tissues, suggesting that TYMP-enhanced TGFβ1 in VSMCs may have a harmful impact and enhance AAA development. However, TYMP does not exhibit a synergistic effect with Ang II in regulating MMP2 production, activation, or TGFβ1 expression. We observed that Ang II does not influence MMP2 production and activity in VSMCs. This finding is consistent with several existing studies that found that Ang II did not affect constitutively expressed MMP2 in VSMCs and cardiac fibroblasts^41, 42^.

In addition to MMP2, uPA is also known to activate TGFβ1. uPA-mediated TGFβ1 activation requires the binding of CD36 and TSP1^43^, a matricellular protein that plays an important role in cell-cell and cell-matrix interaction^17^. Similar to the TGFβ1, although controversial, both salutary^44^ and detrimental^45^ effects of TSP1 have been reported in the AAA milieu. We have extensively studied the role of CD36 in the development of cardiovascular disease and recently found that TSP1-TSR/CD36 signaling enhances VSMC proliferation^46, 47^. CD36 deficiency did not reduce CTGF expression in VSMCs (data not shown), suggesting that TSP1/CD36 signaling does not affect CTGF. Therefore, TSR-induced, TYMP-facilitated CTGF expression is most likely through the activation of the TGFβ1 signaling pathway^17^. CTGF participates in diverse biological processes, including growth and development; however, overexpression of CTGF is correlated with severe fibrotic disorders and a proinflammatory status in VSMCs that leads to endothelial dysfunction^48, 49^. Through multiple positive feedback loops, CTGF could enhance TGFβ1 signaling^48^, further leading to an unbalanced homeostasis of VSMCs environment, which may contribute to the development of AAA. Additional studies are needed to clarify this speculation.

In conclusion, our research uniquely revealed that TYMP, an enzyme in the pyrimidine salvage pathway, possesses multifaceted functions. It plays a significant role in modulating vascular biology, particularly in the functioning of VSMCs and systemic inflammation. Changes in the microenvironment mediated by TYMP possibly contribute to the progression of AAA. This study introduces a new mechanistic target for AAA treatment. Further investigation is required to explore the clinical applicability of this discovery, particularly using the TYMP-selective inhibitor tipiracil, an FDA-approved drug already shown to inhibit thrombosis in mice.

## 5. Acknowledgment

This work is supported by Marshall University Institute Start Fund (to WL), NIH R15HL145573 (to WL), WV-INBRE grant P20GM103434, and West Virginia Clinical and Translational Science Institute Fund supported by the National Institute of General Medical Sciences (U54GM104942). SYC is supported by NIH HL119053 and the Department of Veterans Affairs Merit Review Awards (I01 BX006161). JZ is supported by funds from the Science and Technology Planning Project of Guangdong Province, China (No. 2019B020230003, 2017B090904034, 2017B03031410, 2018B090944002), the National Key Research and Development Program of China (2018YFC1002600), Guangdong Peak Project (DFJH201802), the National Natural Science Foundation of China (No.62006050). HLH is supported by funds from the National Natural Science Foundation of China (82270373) and the Guangdong Basic and Applied Basic Research Foundation (2019B1515120071). The content is solely the responsibility of the authors and does not necessarily represent the official views of the National Institutes of Health and other funding facilities.

We thank Dr. Hong LU for consulting on technical issues in generating the AAA model. We thank Dr. Gang Zhao, Dr. Yueheng Wu, and Dr. Pengju Wen for providing human AAA samples, Dr. Jinsong Huang for providing aorta samples harvested from healthy donors. We thank Dr. Roy L. Silverstein and Dr. Phil Klenotic for providing the TSR peptide.

## 6. Disclosures

None

## Notes

### Competing Interest Statement

The authors have declared no competing interest.

