## Supplemental Materials for "Thymidine Phosphorylase Promotes the Formation of Abdominal Aortic Aneurysm in Mice Fed a Western Diet"

**Short title:** TYMP promotes aneurysm formation.

### These two authors contribute equally to this project.

*** Corresponding author:**

**Wei Li, MD, PhD, FAHA**
Department of Biomedical Sciences, BBSC 241G
Joan C. Edwards School of Medicine at Marshall University
1700 Third Ave, Huntington, WV 25701, USA.


Or

**Jian Zhuang, MD**
Guangdong Provincial People's Hospital, Guangdong Academy of Medical Sciences,
Guangdong Cardiovascular Institute, Guangzhou, China
Address: 106 Zhong Shan Er Lu, Guangzhou, Guangdong, 510080, China


**Supplementary Table 1. AAA patient information.**

| AAA Patient ID | Sex | Age | Race | Primary Diagnosis | Secondary Diagnosis | Complications | AAA diameter (mm) |
|---|---|---|---|---|---|---|---|
| #1 | M | 70 | Asian | AA Dissection | arteriosclerosis | T2DM, hypertension | 53 |
| #2 | M | 56 | Asian | AAA | Arteriosclerosis Obliterans of Lower Extremities | Hyperuricemia, gout, hyperlipoidemia, calculus of gallbladder | 83 |
| #3 | F | 67 | Asian | AAA | coronary atherosclerosis | Hypertension, intervertebral disc herniation | 61 |
| #4 | M | 66 | Asian | AAA | abdominal arteriosclerosis | Hypertension | 50 |
| #5 | M | 58 | Asian | AAA | abdominal arteriosclerosis | Diffuse large B cell lymphoma (DLBCL), bladder stone, hyperplasia of prostate, syphilis | 44 |
| #6 | M | 73 | Asian | AAA | coronary atherosclerosis | Hypertension | 95 |
| #7 | F | 85 | Asian | AAA | coronary atherosclerosis | Anemia, kidney stone, chest effusion | 110 |
| #8 | M | 71 | Asian | AAA | Abdominal atherosclerosis | Hypertension | 80 |
| #9 | M | 84 | White | AAA | aneurysm wall atherosclerosis | Kidney disease, Parkinson’s | information unavailable |
| #10 | F | 91 | White | AAA | abdominal aorta-aneurysm repair. changes with aneurysm wall atherosclerosis | Neck and lumbar surgery, pacemaker/defibrillator | information unavailable |

| | | | | | | | |
|---|---|---|---|---|---|---|---|
| #11 | F | 75 | White | AAA | atherosclerosis consistent with aneurysm | VP shunt, hydrocephalus | information unavailable |
| #12 | M | information unavailable | White | AAA | plaque; aneurysm repair; atherosclerotic plaque | information unavailable | information unavailable |
| #13 | M | 69 | White | AAA | complicated atherosclerosis | emphysema/COPD, pacemaker | information unavailable |
| #14 | M | 77 | White | AAA | consistent with dissecting aneurysm | information unavailable | information unavailable |
| #15 | M | 87 | White | AAA | focal cystic medial degeneration | Arthritis, degenerative disk disease | information unavailable |
| #16 | M | information unavailable | White | AAA | calcification | information unavailable | information unavailable |

**Supplementary Table 2. Primers for testing the targeted genes by real-time PCR.**

| Targets | Targeting Species | Forward Sequence (5'-3') | Reverse Sequence (5'-3') |
|---|---|---|---|
| TYMP | human | TGATCCGCATGAAGCGAGAC | CAGATCCATGCCCCGAAGT |
| GAPDH | human | AATCCCATCACCATCTTCCAG | GAGCCCCAGCCTTCTCCAT |
| TIMP-2 (Pair#1) | Rat | GCATCACCCAGAAGAAGAGC | GTTCAAGAAACGGACGTAGT |
| TIMP-2 (Pair #2) | Rat | CAAGTTCTTTGCCTGCATCA | GAACTGTAGGGAAGGACCT |
| MMP-2 (Pair#1) | Rat | TCCCCTGATGCTGATACTGAC | TTTTCTAACTACGGCACATGC |
| MMP-2 (Pair#2) | Rat | GCTGTGGACTCTAGGAGAAGGA | AAGGTGGTGCATGTTGAAACTC |
| TGF β1 | Rat | CGCCTGCAGAGATTCAAGTC | TGACGTCAAAAGACAGCCAC |
| GAPDH | Rat | CAGAACATCATCCCTGCATC | CTGCTTCACCACCTTCTTGA |
| MMP2 | Mouse | CCCGATCTACACCTACACCAA | AAACCGGTCCTTGAAGAAGAA |
| TYMP | Mouse | CGCGGTGATAGATGGAAGAGCA | CCGCTGATCATTGGCACCTTAC |
| GAPDH | Mouse | CAGCAACTCCCACTCTTCCACCTTCG | GGCCTCTCTTGCTCAGTGTCCTTGCT |

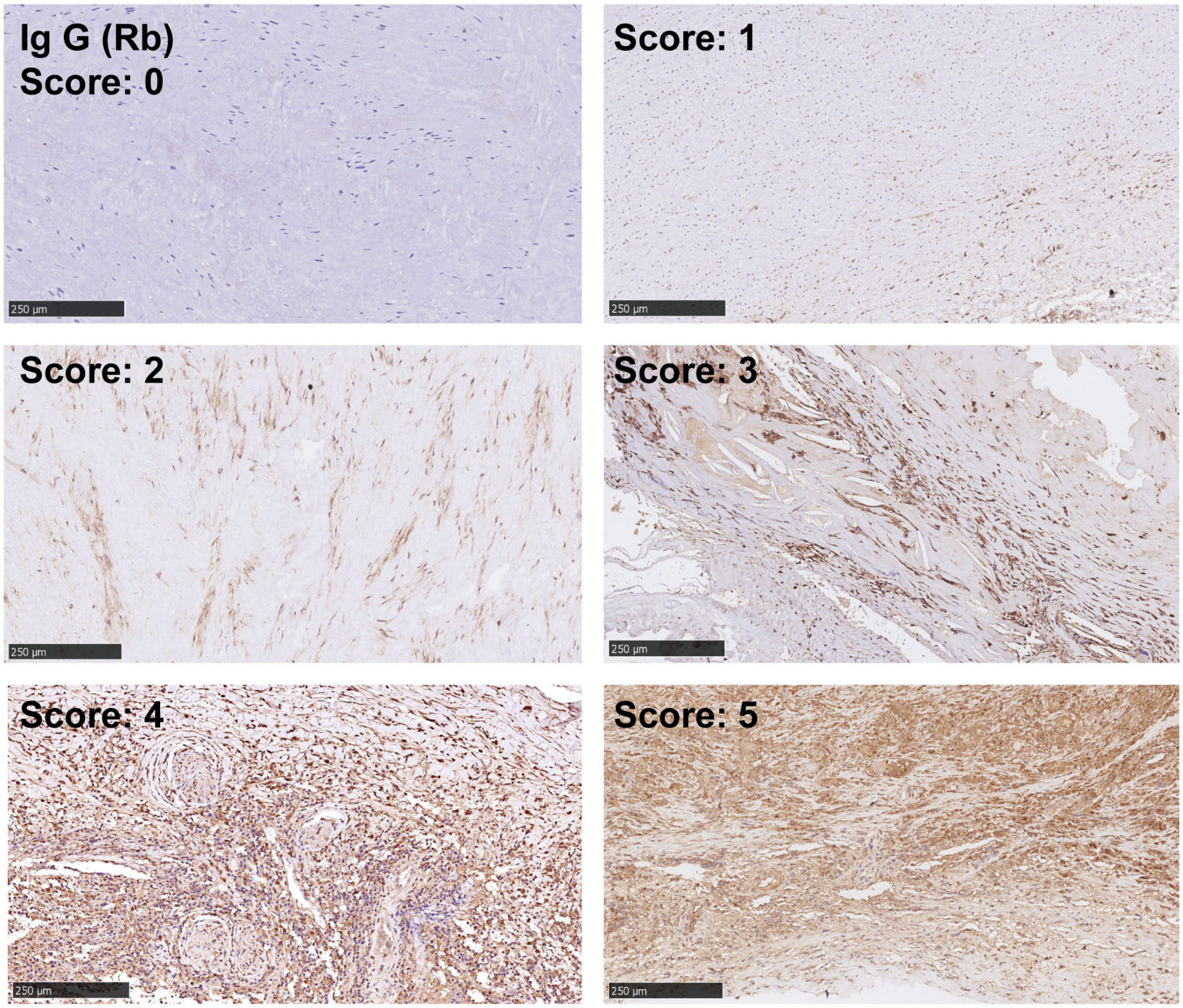


**Supplementary Figure 1.** Histochemical staining images were scored based on staining intensity. We adopted a 6-point scale system, and representative images with score assignments are provided.

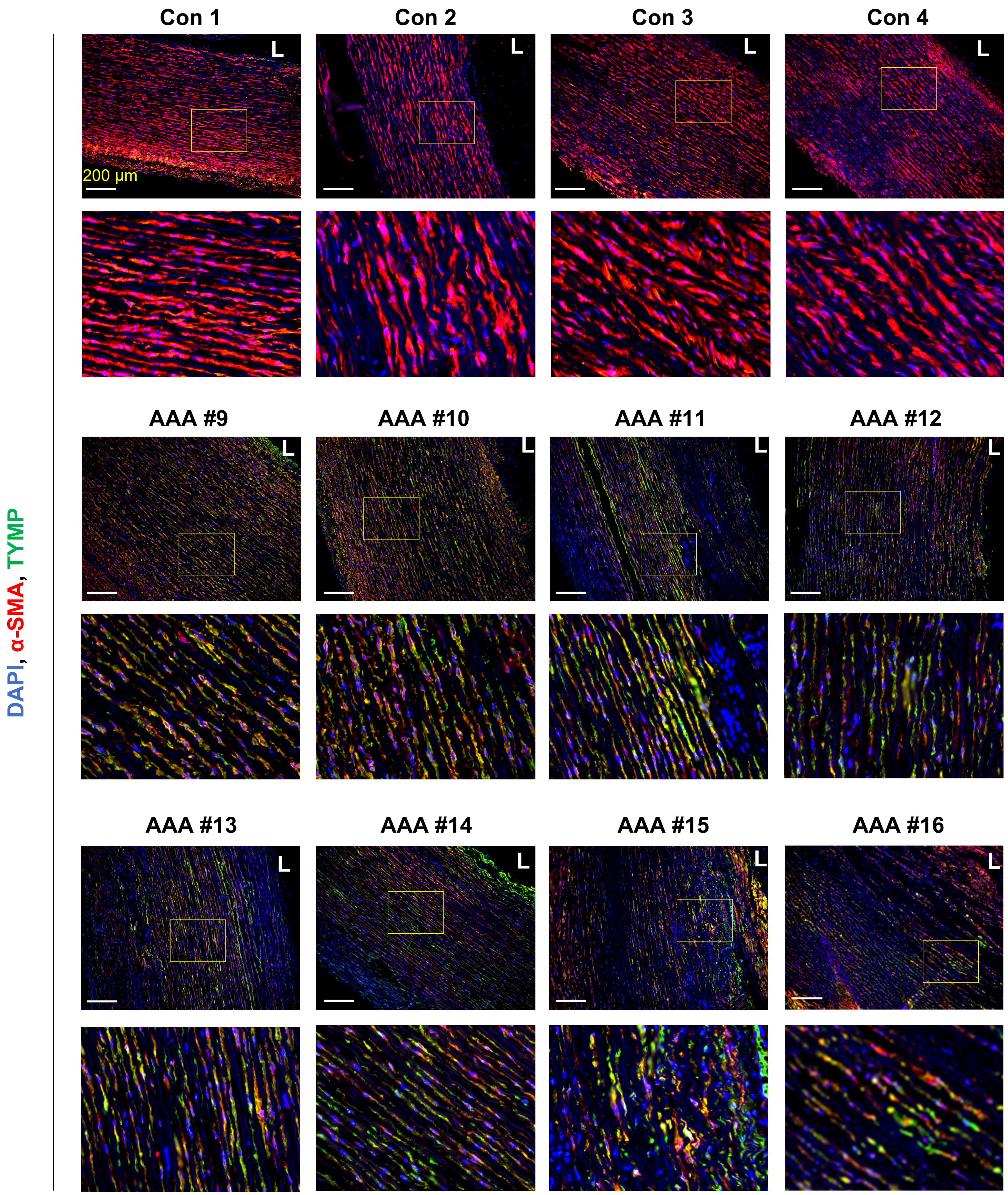


**Supplementary Figure 2. TYMP expression is increased in human AAA vessel wall.** Human AAA vessel wall and healthy control aorta vessel wall (Con) were sectioned and double stained for TYMP and α-SMA. Nuclei were stained with DAPI.

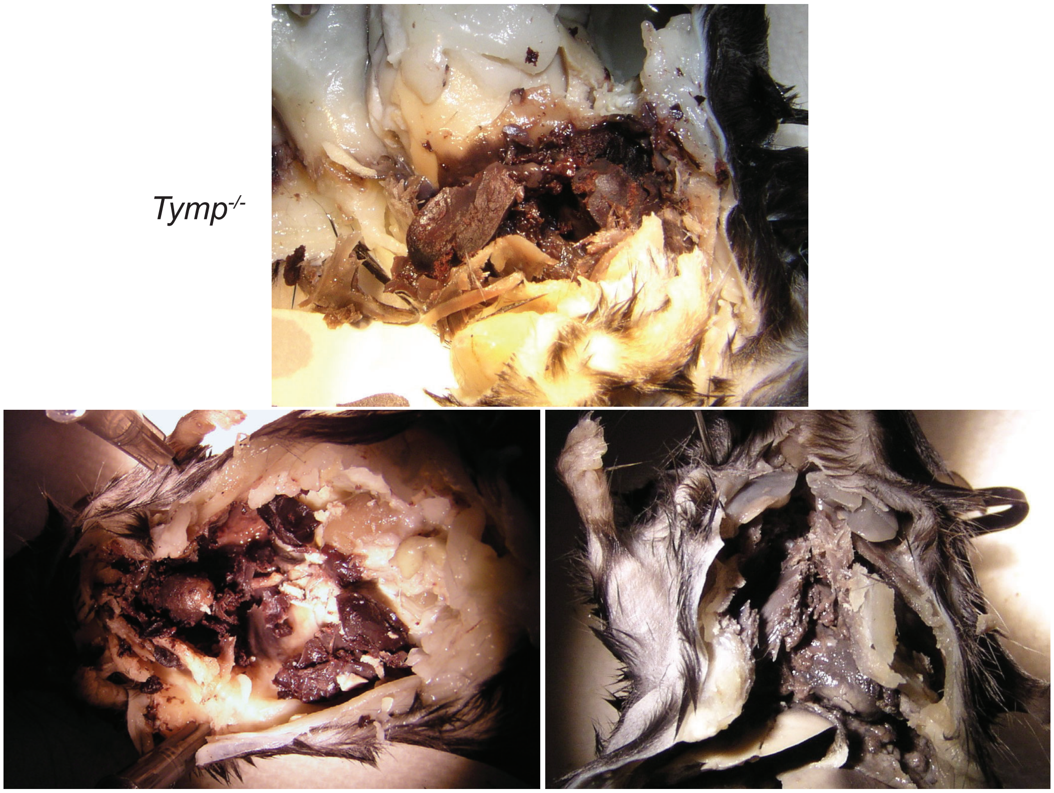


**Supplementary Figure 3.** Necropsy of mice died after Ang II perfusion.

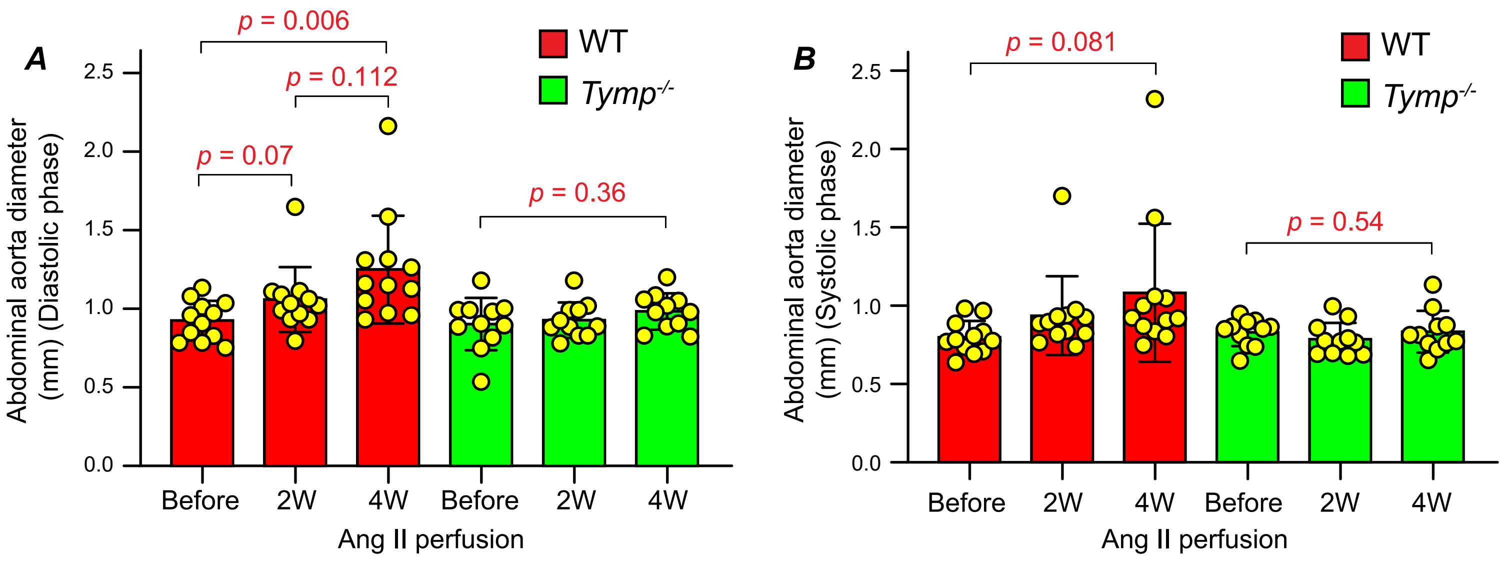


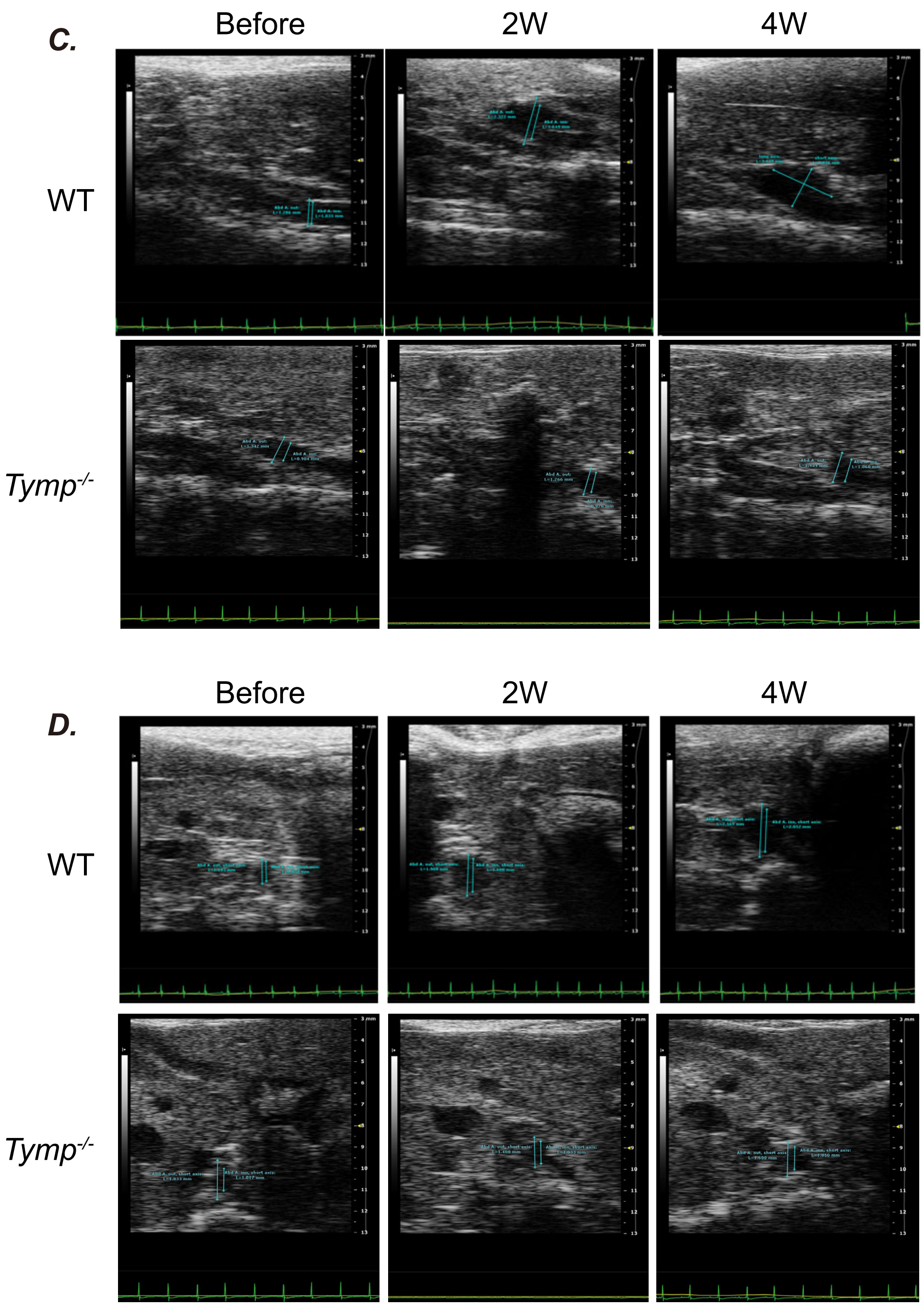


**Supplementary Figure 4. Thymidine phosphorylase (TYMP) deficiency reduced the prevalence of abdominal aortic aneurysm formation (AAA) in mice.** Mice were treated as mentioned in Fig. 2. ***A.*** Inner diameter of abdominal aorta in the diastolic phase. ***B.*** Inner diameter of abdominal aorta in the systolic phase. ***C.*** Representative echo images in the diastolic phase. ***D.*** Representative echo images in the systolic phase.

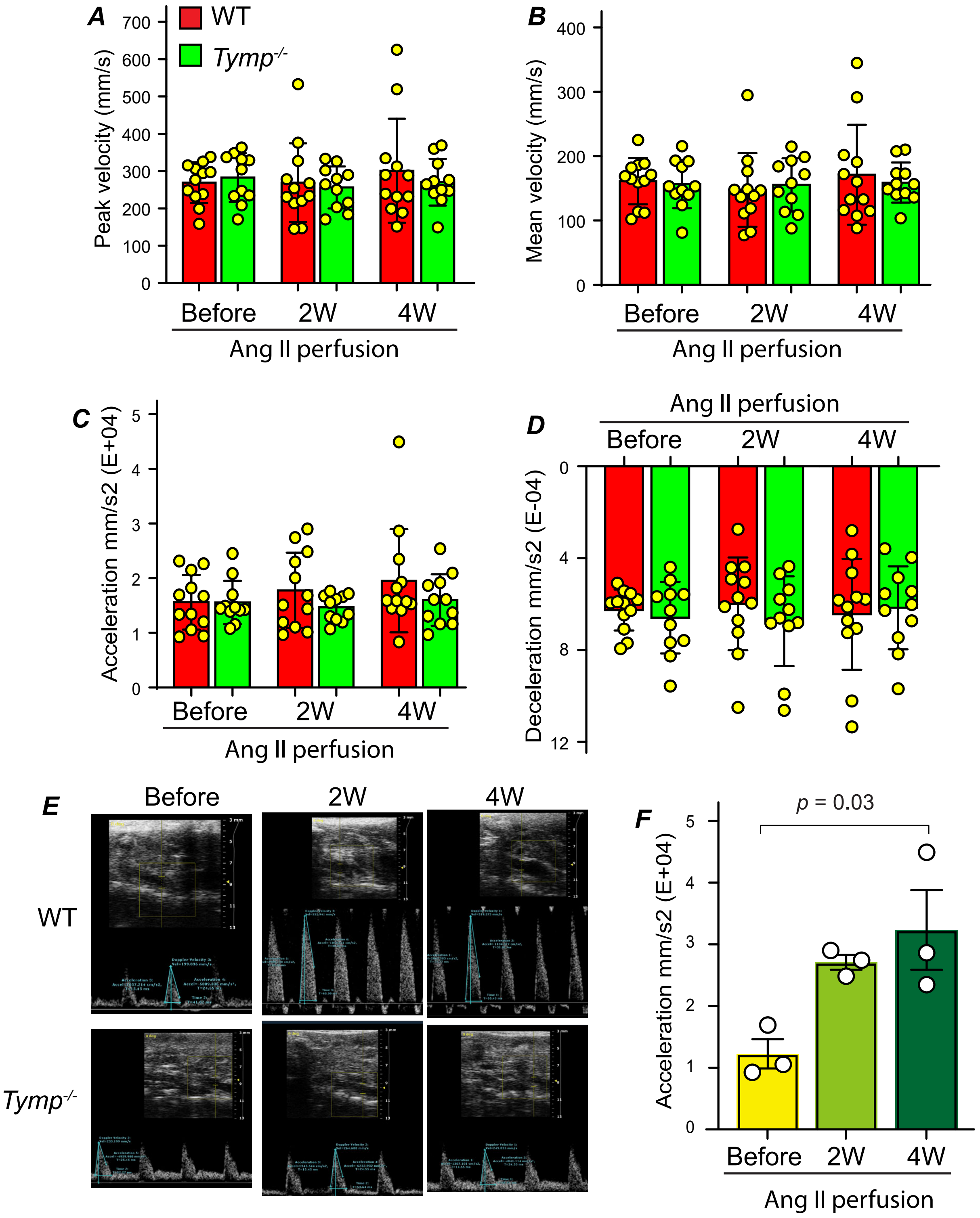


**Supplementary Figure 5. Thymidine phosphorylase (TYMP) deficiency reduced the prevalence of abdominal aortic aneurysm formation (AAA) in mice.** Mice were treated as mentioned in Fig. 2. ***A.*** Peak velocity, ***B.*** mean velocity, ***C.*** acceleration and ***D.*** deacceleration were determined. ***E.*** Representative acceleration echo images in the WT and *Tymp*$^{-/-}$ mice. ***F.*** Statistical analysis of acceleration in the three WT mice with AAA.

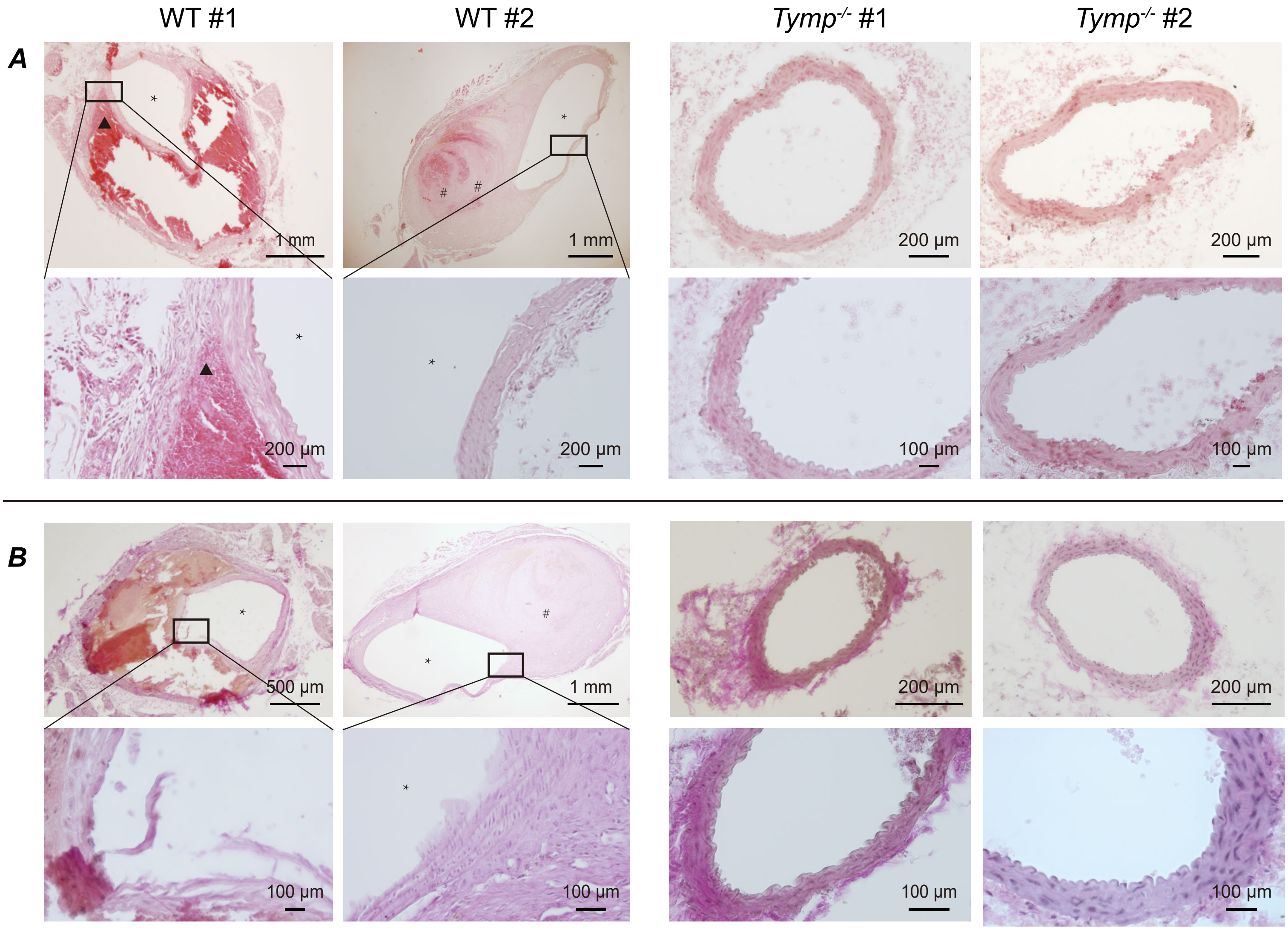


**Supplementary Figure 6. TYMP deficiency attenuates distortion of vessel wall structure in the mouse AAA model.** ***A.*** H&E staining. ***B.*** Elastica van Gieson (EVG) staining. * indicates vessel true lumen. # indicates hematoma.

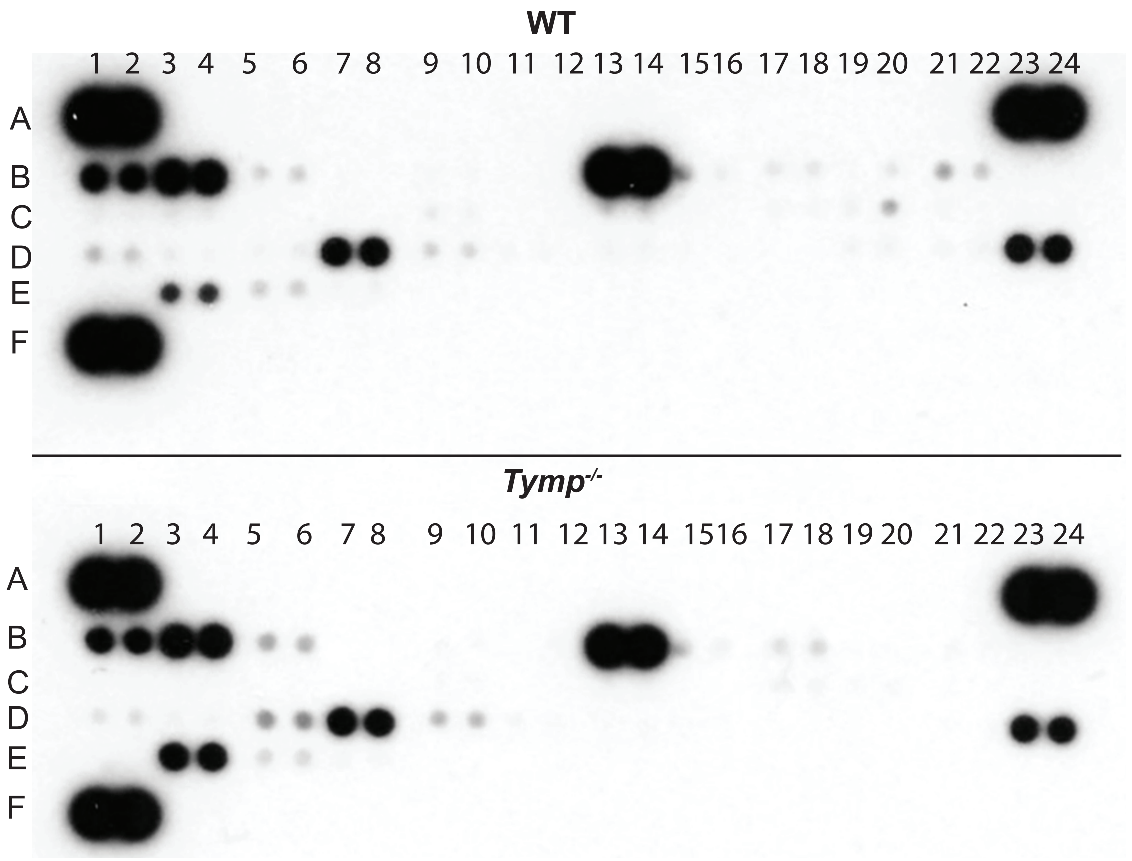


**Supplementary Figure 7. Expression of plasma cytokines in WT and *Tymp*$^{-/-}$ mice fed a Western diet.** Plasma pooled from 6 WT mice (including 3 mice with AAA and 3 randomly selected) and 6 *Tymp*$^{-/-}$ mice (including 1 mouse with AAA and 5 randomly selected) were subjected to the Proteome Profiler Mouse Cytokine Array, Panel A.

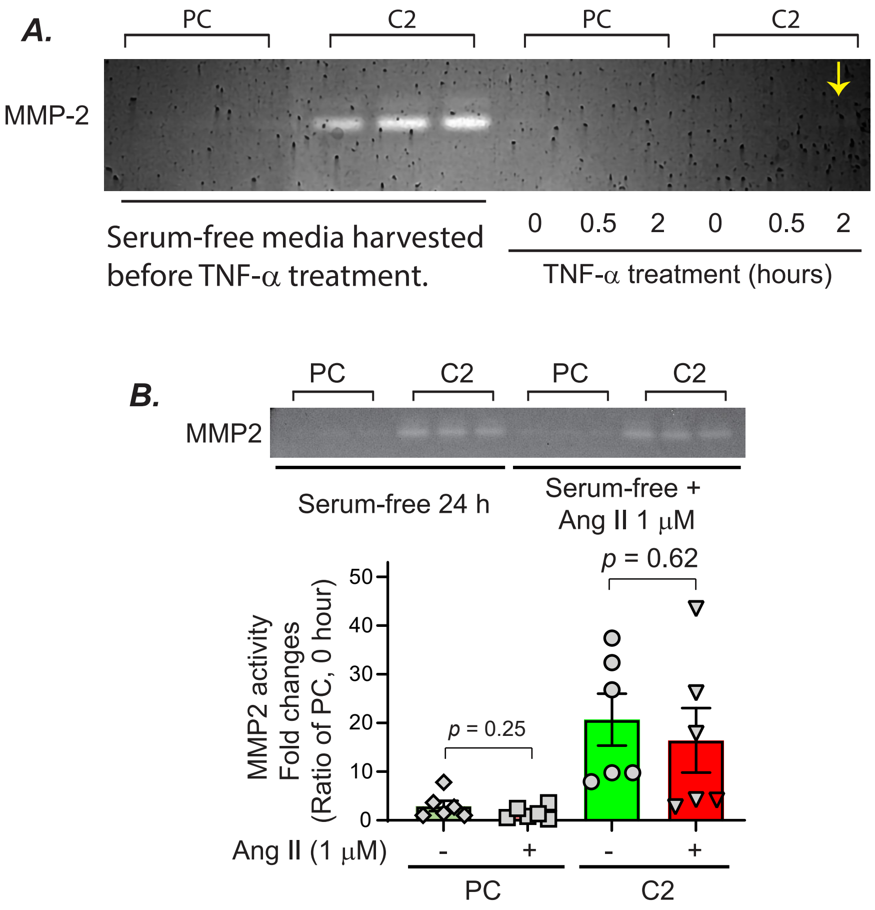


**Supplementary Figure 8. TYMP enhances MMP2 production and secretion in VSMCs.** ***A.*** C2 and PC cells were synchronized by serum starvation overnight, and then treated with TNF-α (10 ng/mL) for the indicated durations. Both the ovenight-conditioned media and media after TNF-α treatment was subjected to Zymography. The arrow indicates the weak MMP2 band. **B.** PC and C2 cells were serum-starved for 24 hours and then treated with Ang II at a final concentration of 1 μM in serum-free DMEM for additional 24 hours. The culture media were collected and used for gelatin zymography.

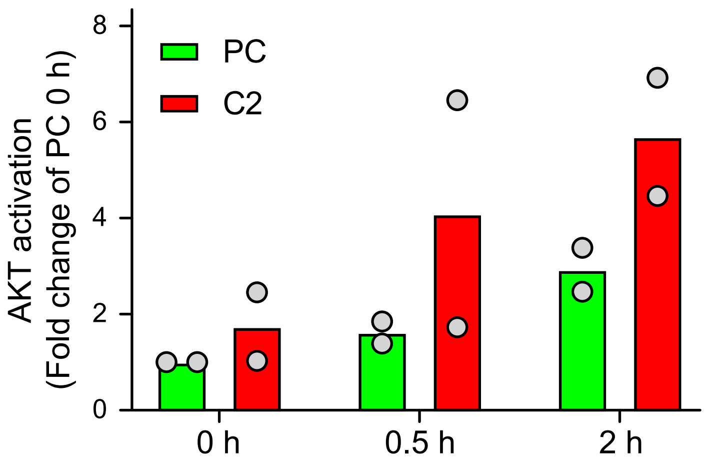


**Supplementary Figure 9. TYMP overexpression enhances AKT signaling activation in VSMCs.** Serum starvation synchronized C2 and PC cells were treated with TNF-α for the indicated durations, and cell lysates were used for Western blot assay of AKT activation as shown in Figure 6C. Band intensity was analyzed using ImageJ, and p-AKT expression was adjusted with the expression of total AKT. Data were shown as fold change of PC at 0 h.

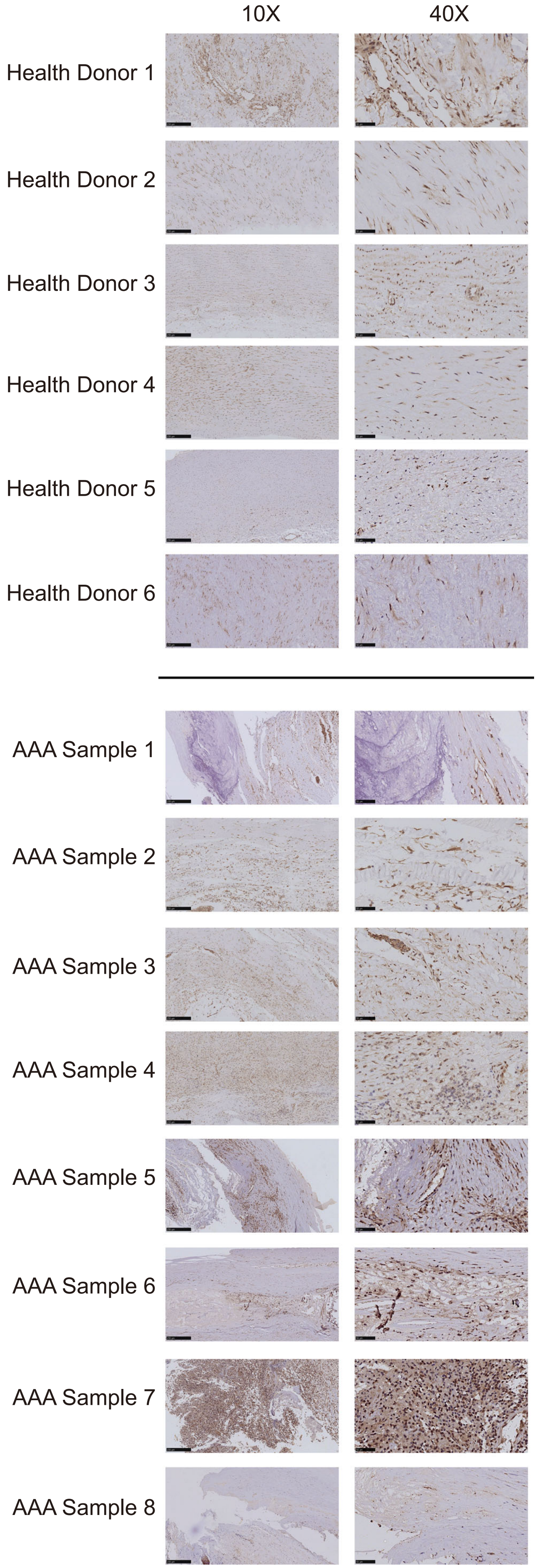


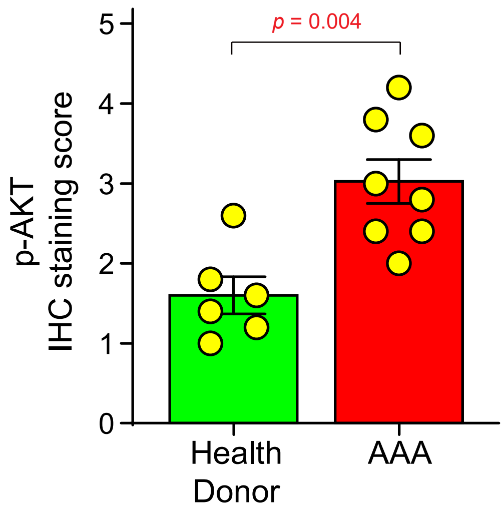


**Supplementary Figure 10. Evaluation of AKT activation in healthy human aortic vessel wall and AAA samples.** Parafin-embedded human aortic tissues were sent to Bios Biological (https://biossci.com/), a professional company for histological study. Immunohistochemical staining of p-AKT was conducted using an antibody against phospho-pan-AKT1/2/3 (Ser473) (AF0016, Affinity), which detects endogenous levels of pan-AKT1/2/3 only when phosphorylated at Sersine 473. Positive staining is indicated by brown coloration, while nuclei were staining with hematoxylin (blue). The images were scanned and scored by lay individuals based on staining intensity showed in Supplementary Figure 1, and score data were used for statistical analysis, with results presented in bar graph format.

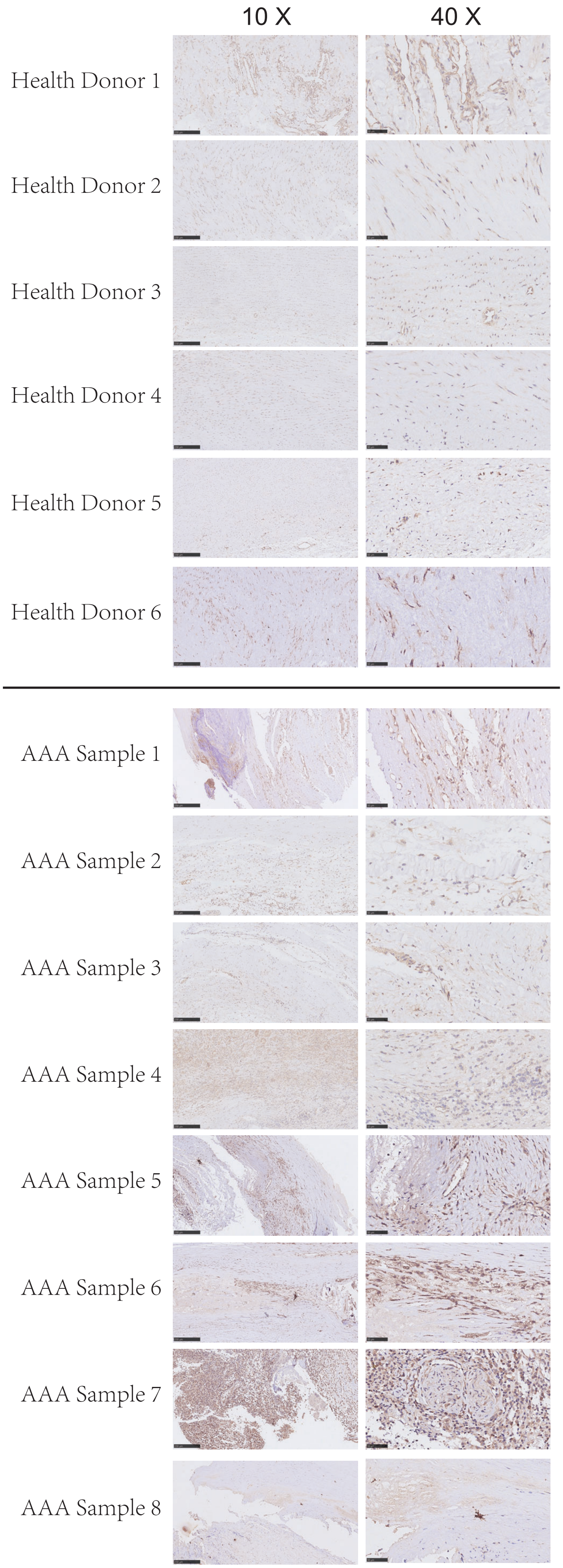


**Supplementary Figure 11.** IHC of TGFβ1 in human healthy donor aorta or in the AAA vessel walls. Brown indicates positive staining.

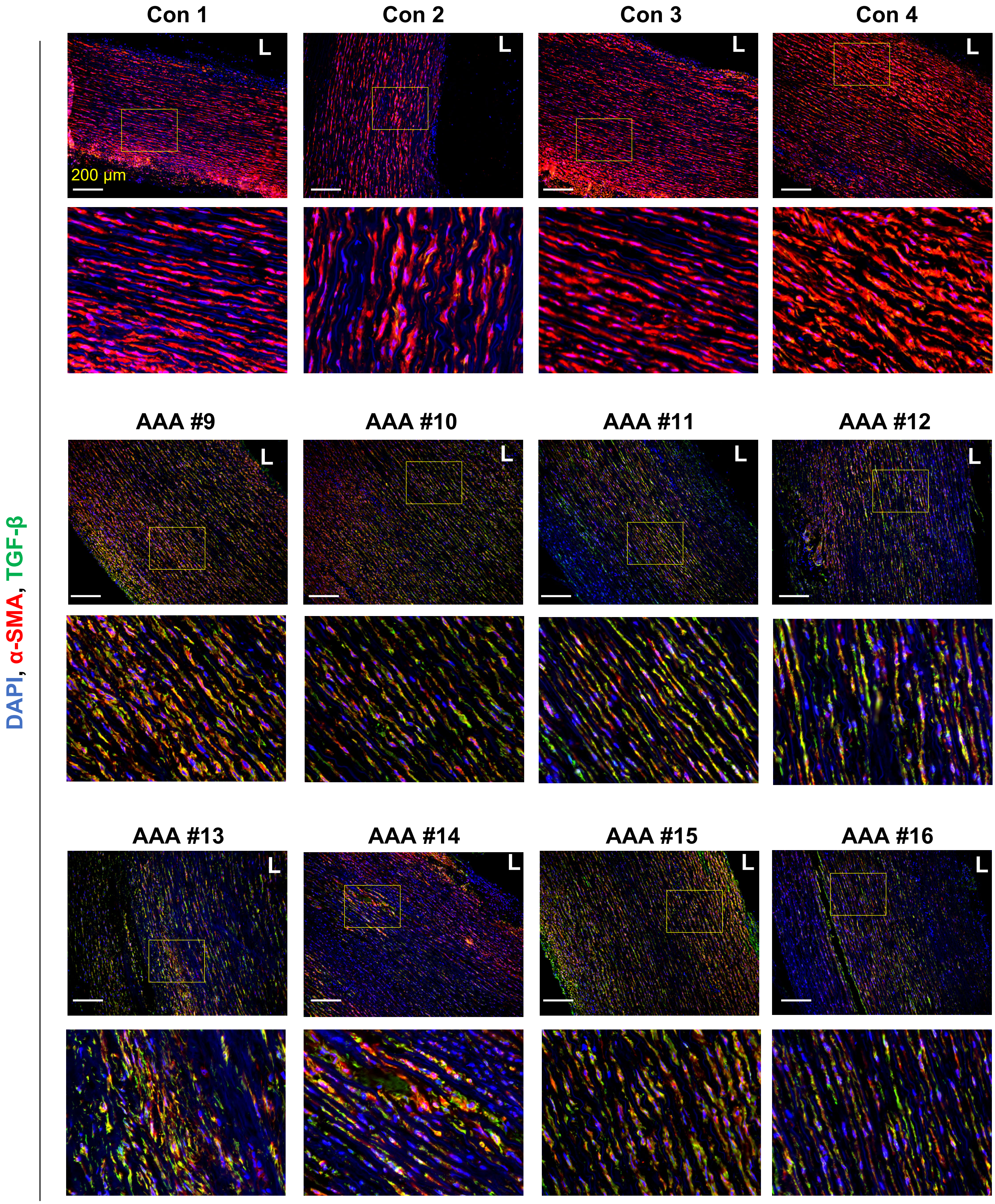


**Supplementary Figure 12. TGFβ1 expression is increased in human AAA vessel wall.** Human AAA vessel wall and healthy control aorta vessel wall (Con) were sectioned and double stained for TGFβ1 and α-SMA. Nuclei were stained with DAPI.

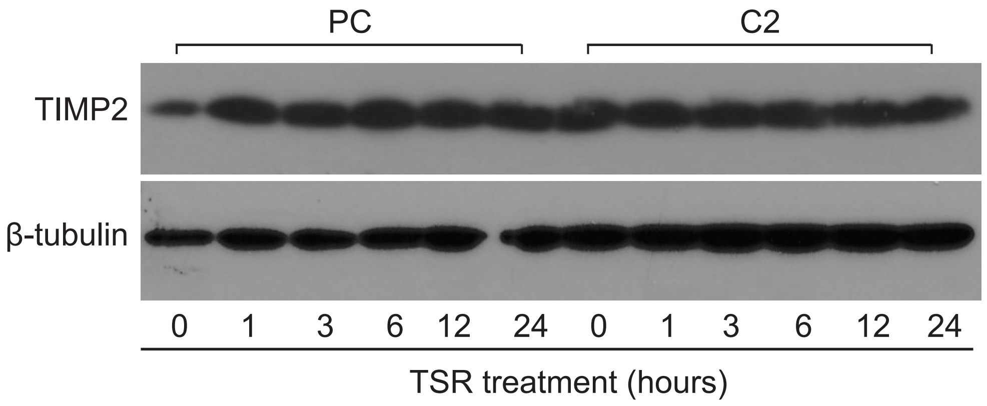


**Supplementary Figure 13. TYMP overexpression does not affect TIMP2 expression in VSMCs.** As mentioned in Fig. 7H, serum-starved PC and C2 cells were treated with serum-free media in the presence of 10 μg/mL TSR for the indicated durations. TIMP2 levels were examined by Western blot analysis, with β-tubulin blotted as loading control.
